# Increased core body temperature exacerbates defective protein prenylation in mouse avatars of mevalonate kinase deficiency

**DOI:** 10.1101/2022.02.28.480959

**Authors:** Marcia A. Munoz, Oliver P. Skinner, Etienne Masle-Farquhar, Julie Jurczyluk, Ya Xiao, Emma Fletcher, Esther Kristianto, Mark P. Hodson, Seán I. O’Donoghue, Sandeep Kaur, Robert Brink, David Zahra, Elissa K. Deenick, Kristen Perry, Avril A.B. Robertson, Sam Mehr, Pravin Hissaria, Catharina M. Mulders-Manders, Anna Simon, Michael J. Rogers

**Affiliations:** Garvan Institute of Medical Research and School of Clinical Medicine, UNSW Sydney, Sydney, NSW, Australia; Victor Chang Cardiac Innovation Centre, Victor Chang Cardiac Research Institute, Sydney, Australia; School of Pharmacy, University of Queensland, Woolloongabba, Australia; School of Chemistry and Molecular Biosciences, University of Queensland, Brisbane, Australia; Royal Children’s Hospital, Melbourne, Australia; Royal Adelaide Hospital, SA Pathology and University of Adelaide, Adelaide, Australia; Department of Internal Medicine, Radboudumc Expertise Centre for Immunodeficiency and Autoinflammation, Radboud University Medical Centre, Nijmegen, The Netherlands

**Keywords:** prenylation, autoinflammatory, mevalonate kinase deficiency, HIDS, Rab GTPase, IL-1β, NLRP3, mevalonic acid, thermogenesis

## Abstract

Mevalonate kinase deficiency (MKD) is caused by biallelic loss-of-function mutations in *MVK*, leading to recurrent fevers and systemic inflammation. We describe new mouse avatars of MKD bearing p.Val377Ile (the commonest variant) or deletions in *Mvk*. Compound heterozygous mice recapitulated the biochemical phenotype of MKD, with build-up of unprenylated GTPases and increased plasma mevalonic acid. Mice with different deficiencies in mevalonate kinase revealed new insights into the genotype-phenotype relationship and mirrored the variability in the prenylation defect in human MKD, with p.V377I homozygous mice having a milder phenotype than compound heterozygous animals. The inflammatory response to LPS was enhanced in compound heterozygous mice *in vivo* and elevated serum interleukin-1β was abrogated by NLRP3 inflammasome inhibition. Increased temperature dramatically but reversibly exacerbated the deficit in the mevalonate pathway and defective prenylation *in vitro* and *in vivo*, highlighting increased body temperature as a likely trigger of inflammatory flares and an additional potential target for future therapeutic approaches.

## INTRODUCTION

The mevalonate pathway is an essential, ubiquitous metabolic pathway required for the biosynthesis of long-chain isoprenoid lipids such as farnesyl diphosphate and geranylgeranyl diphosphate (GGPP). The latter are the building blocks of numerous other metabolites, including sterols, but are also necessary for the post-translational prenylation of >300 proteins, particularly small GTPases such as those of the Rho/Rac/Rap and Rab superfamily (Maurer-Stroh et al., 2007; Wang and Casey, 2016). Dysregulation of the mevalonate pathway has been associated with a wide variety of human diseases, and particularly with inflammation. The most striking example of this is the genetic autoinflammatory disorder mevalonate kinase deficiency (MKD). MKD is an inborn error of metabolism caused by autosomal recessive inheritance of mutations in the *MVK* gene (Drenth et al., 1999; Houten et al., 1999) encoding mevalonate kinase (MK; ATP:(R)-mevalonate 5-phosphotransferase, EC 2.7.1.36), a proximal enzyme in the mevalonate pathway (Brennenstuhl et al., 2021; Houten et al., 2000). Understanding of the molecular mechanisms underpinning MKD pathogenesis remain largely unknown due to the lack of appropriate genetic tools to study this disease.

MKD encompasses a severe manifestation - mevalonic aciduria (OMIM 610377), and a milder form - a periodic fever syndrome known as hyperimmunoglobulinemia D syndrome (HIDS, OMIM 260920) (Mandey et al., 2006b; Simon et al., 2004b). In HIDS, the inflammatory symptoms usually appear in early childhood and feature regular episodes of high fever and systemic inflammation separated by intervals of normal health. The most frequent pathogenic variant in HIDS is a G>A missense mutation in exon 11 of *MVK*, resulting in a p.Val377Ile substitution (Boursier et al., 2021; Houten et al., 2000; Simon et al., 2004b; Ter Haar et al., 2016). Individuals homozygous or compound heterozygous for *MVK*^*V377I*^ have 1-20% residual mevalonate kinase (MK) enzyme activity (Cuisset et al., 2001; Frenkel et al., 2001), but why such a conservative amino acid substitution affects the MK enzyme remains unknown. Mevalonic aciduria is caused by homozygous or compound heterozygous *MVK* variants that affect enzyme function or expression more severely than *MVK*^*V377I*^ (Brennenstuhl et al., 2021), and thus is associated with extremely low (<0.5%) or undetectable residual MK activity (Hoffmann et al., 1986; Hoffmann et al., 1993; Poll-The et al., 2000). In contrast to HIDS, mevalonic aciduria patients present with persistent systemic inflammation (Simon et al., 2004b; Ter Haar et al., 2016), as well as neurologic impairment (including cerebellar atrophy, dystonia and ataxia). The build-up of the substrate of MK, mevalonic acid (MA, or the lactone derivative), in urine and plasma is a characteristic biochemical feature of mevalonic aciduria and also occurs during inflammatory flares in HIDS (Frenkel et al., 2001; Haas and Hoffmann, 2006; Hoffmann et al., 1986; Hoffmann et al., 1993; Poll-The et al., 2000).

Another hallmark of MKD is elevated serum levels of IL-1β and increased release of IL-1β from PBMCs and fibroblasts of patients (Drenth et al., 1995; Normand et al., 2009). This enhanced IL-1β production is generally attributed to increased activation of the multi-protein complexes (inflammasomes) responsible for proteolytic processing of proIL-1β into its mature form (Chan and Schroder, 2020). Furthermore, we recently provided the first direct evidence that prenylation of Rab and Rap1A GTPases is deficient in MKD PBMCs, and found that accumulation of unprenylated Rab proteins could be a diagnostic indicator of MKD (Munoz et al., 2017; Munoz et al., 2019). Still, many questions remain unanswered. How loss of MK activity and disruption of the mevalonate pathway lead to enhanced inflammasome activity is unclear but appears to involve loss of synthesis of the isoprenoid lipid GGPP (Frenkel et al., 2002; Mandey et al., 2006a) necessary for protein prenylation. Furthermore, the identity of the inflammasome(s) affected by defective protein prenylation is still controversial, with evidence for enhanced activation of NLRP3 (Skinner et al., 2019), as well as pyrin (Akula et al., 2016; Park et al., 2016). This remains a critical question, given the immense potential of inflammasome inhibitors as new therapeutic approaches for autoinflammatory disorders such as MKD. Also, inflammatory flares in MKD are commonly triggered by stress, strenuous exercise, vaccinations or infection (Drenth et al., 1994; Frenkel et al., 2001; Govindaraj et al., 2020; Houten et al., 2000) but the underlying mechanisms remain unknown. An interesting proposal is that mutations in *MVK* may lead to temperature-sensitive changes in folding or stability of the enzyme (Houten et al., 2002; Mandey et al., 2006b). Hence, elevations in body temperature could potentially further compromise MK activity and exacerbate the underlying defect in protein prenylation. This hypothesis, however, remains to be tested *in vivo* in a relevant physiological model of MKD.

A major reason for our poor understanding of the mechanisms of disease in MKD is the lack of genetic animal models, since complete loss of *Mvk* expression is lethal in homozygous *Mvk* knockout mice (Hager et al., 2007). One study used conditional knockout of a prenyltransferase gene to prevent protein prenylation (Akula et al., 2016). Other *in vitro* and *in vivo* approaches utilised pharmacologic inhibitors such as statins, bisphosphonates and prenyltransferase inhibitors. These compounds, although blocking different steps of the mevalonate pathway, act commonly to inhibit downstream protein prenylation (Rogers et al., 2020) and thus, have been used as models of MKD (Frenkel et al., 2002; Jurczyluk et al., 2016; Kuijk et al., 2008a; Mandey et al., 2006a; Marcuzzi et al., 2008; Marcuzzi et al., 2013; Massonnet et al., 2009; Montero et al., 2000; Skinner et al., 2019). However, whether any of these pharmacological or genetic models accurately mimic the defect in prenylation that occurs in MKD is not known. Another important limitation of these models is that they do not permit studies on complex physiological mechanisms that may affect the mutant MK enzyme and thus contribute to inflammatory flares. To fill this fundamental knowledge gap and better understand the link between defective protein prenylation, excessive inflammation and inflammatory flares, we used CRISPR/Cas9 gene editing to create physiologically-relevant mouse models of MKD carrying bi-allelic mutations in *Mvk*, including the commonest pathogenic variant p.V377I.

## RESULTS

### *Mvk* mutant mice have reduced MK activity

Using CRISPR/Cas9 gene editing of C57BL/6J mouse embryos we created 3 mouse lines carrying heterozygous deletions of 8bp (*Δ8*), 13bp (*Δ13*) or 91bp (*Δ91*) in exon 11 (Suppl. Fig 1). The *Δ8* mutation resulted in a frame shift following the codon for T370 and a predicted extension of 24 amino acids at the C terminus; the *Δ13* mutation caused a frameshift after the codon for A734 and premature termination of the C-terminus (Suppl. Fig. 1). The *Δ91* frameshift mutation was predicted to cause loss of residues L348-S378, replaced by an altered sequence of 7 amino acids (residues 348-354, LEQPEVE > PHTQLQL) and deletion of all remaining residues from 355 onwards (Suppl. Fig. 1).

*Mvk* mutant mice were intercrossed to breed animals homozygous for *Mvk*^*Val377Ile*^ (hereafter abbreviated to *Mvk*^*VI/VI*^ or simply VI/VI). Compound heterozygous mice were generated with the biallelic mutations *Mvk*^*VI*^ and either *Δ8, Δ13* or *Δ91* (i.e. *Mvk*^*VI/Δ8*^, *Mvk*^*VI/Δ13*^ or *Mvk*^*VI/Δ91*^, respectively). *Mvk*^*VI/VI*^, *Mvk*^*VI/Δ8*^, *Mvk*^*VI/Δ13*^ and *Mvk*^*VI/Δ91*^ mice were viable, born in the expected Mendelian ratio, and aside from a slightly lower (but not significantly different) mean body weight of *Mvk*^*VI/Δ91*^ 8 week male and female mice, these mutant animals did not differ in development or appearance from wildtype or littermate heterozygous mice (Fig. 1A,B). No homozygous offspring with *Δ8, Δ13* or *Δ91* mutations in exon 11 were ever born, indicating that these were amorphic mutations. No C-terminal truncated form of MK of the predicted mass could be detected by western blot analysis of liver (which has extremely high levels of MK) from *Δ91* mice, using an antibody that binds to the mid-region of MK (Suppl. Fig. 2), suggesting lack of expression or increased degradation of the mutant *Δ91* MK protein.

**Fig. 1.**
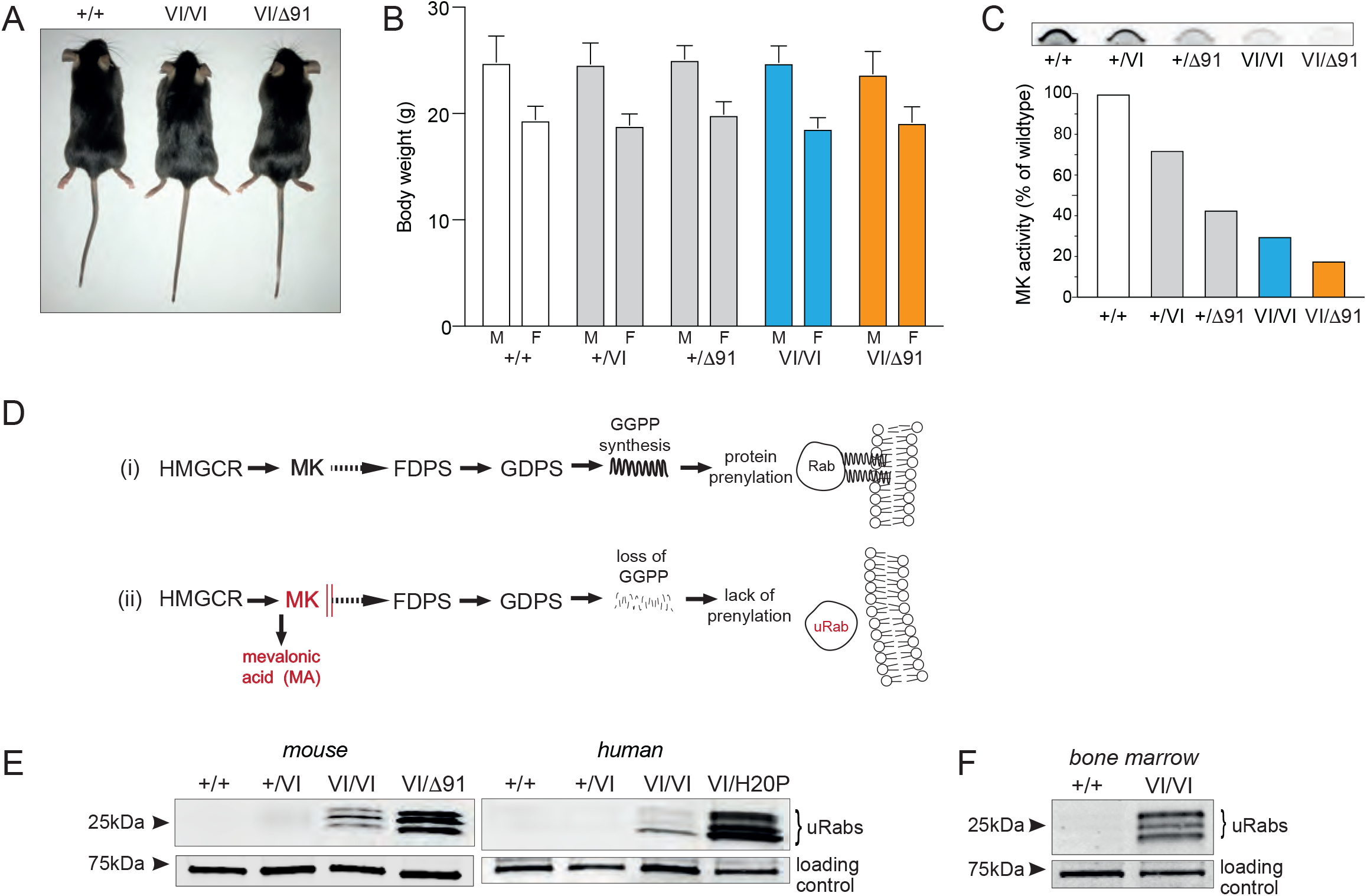
*Mvk*^*VI/VI*^ and *Mvk*^*VI/Δ91*^ mice recapitulate the defective prenylation phenotype of mevalonate kinase deficiency in humans. A,B) Appearance and body weight of wildtype and *Mvk* mutant male (M, n=12) and female (F, n=11) mice (mean ± SD). Mice shown in A) were 12-week-old males. C) Mevalonate kinase (MK) activity in liver homogenates, expressed as a percentage of activity in wildtype liver (values representative of measurements in two mice per genotype). D) Diagram of the mevalonate pathway leading normally (i) to protein prenylation (HMGCR, HMG-CoA reductase; MK, mevalonate kinase; FDPS, farnesyl diphosphate synthase; GDPS, geranylgeranyl diphosphate synthase; lack of MK activity (ii) leads to build-up of mevalonic aid (MA), loss of geranylgeranyldiphosphate (GGPP) synthesis and accumulation of unprenylated Rab (uRab) proteins. E) Comparison of uRabs in spleen cells from *Mvk*^*VI/VI*^ (VI/VI) and *Mvk*^*VI/Δ91*^ (VI/Δ91) mice and in PBMCs from MKD patients (genotypes *MVK*^*VI/VI*^ and *MVK*^*VI/H20P*^). Wildtype and heterozygous genotypes were used as mouse and human controls, and an endogenous, biotinylated 73kDa protein as loading control. F) Detection of uRabs in bone marrow from homozygous *Mvk*^*VI/VI*^ (VI/VI) mice.

Consistent with the *Δ91* mutation causing complete loss of function, heterozygous *Mvk*^*+/Δ91*^ mice had approximately 50% normal MK activity in liver homogenates compared to wildtype counterparts (Fig. 1C). Furthermore, animals carrying the milder p.V377I mutation in one allele (*Mvk*^*+/VI*^) had 73% residual MK activity, whereas *Mvk*^*VI/VI*^ homozygous mice had 19% residual activity. Importantly, *Mvk*^*VI/Δ91*^ mice had the lowest residual MK activity (9% compared to wildtype) (Fig. 1C). Similar results were obtained with bone marrow extracts (18% and 5% residual MK activity in *Mvk*^*VI/VI*^ and *Mvk*^*VI/Δ91*^ bone marrow, respectively, compared to wildtype). Together, this suggests that a single *Mvk*^*VI*^ allele conferred approximately 10% residual MK activity *in vivo*.

### Mutations in mevalonate kinase cause a similar pattern of defective protein prenylation in immune cells in mice and humans

In humans, mutations in *MVK* that reduce MK activity lead to decreased synthesis of isoprenoid lipids via the mevalonate pathway (Fig. 1D), thereby causing a deficiency in protein prenylation (Jurczyluk et al., 2016; Munoz et al., 2017; Munoz et al., 2019). We analysed immune cell populations from *Mvk* mutant mice for evidence of defective protein prenylation, using an *in vitro* prenylation assay (Ali et al., 2015; Jurczyluk et al., 2016; Rogers et al., 2020) to measure the buildup of unprenylated Rab GTPases (uRabs). Unprenylated forms of small GTPases do not accumulate in cells under normal conditions and, like wildtype mice, heterozygous *Mvk*^*+/VI*^, *Mvk*^*+/Δ8*^, *Mvk*^*+/Δ13*^ and *Mvk*^*+/Δ91*^ animals had barely detectable levels of uRabs in spleen cells or bone marrow (Fig. 1E, Suppl. Fig. 3A-C). Homozygous *Mvk*^*VI/VI*^ mice had a mild accumulation of uRabs (a cluster of bands of molecular mass 23-27kDa) in spleen cells and bone marrow cells (Fig. 1E, Suppl. Fig. 3A). This mild prenylation defect was more obvious when the sensitivity of the *in vitro* Rab prenylation assay was increased by lengthening the duration of the incubation with Rab GGTase/REP-1 (with *Mvk*^*VI/VI*^ bone marrow, Fig. 1F). By contrast, cells from compound heterozygous *Mvk*^*VI/Δ91*^ mice showed a clear and substantial accumulation of uRabs (spleen, Fig. 1E; bone marrow and peripheral blood mononuclear cells/PBMCs, Suppl. Fig. 3A,D) to levels almost 20 times higher than in *Mvk*^*VI/VI*^ or heterozygous or wildtype mice (Suppl. Fig. 3B). *Mvk*^*VI/Δ91*^ bone marrow cells also showed a clear accumulation of unprenylated Rap1A (uRap1A) detectable by western blotting (Suppl. Fig. 3A). Similar levels of unprenylated uRabs and uRap1A were found in cells from compound heterozygous *Mvk*^*VI/Δ8*^ and *Mvk*^*VI/Δ13*^ animals (Suppl. Fig. 3A,B). We therefore chose to focus on *Mvk*^*VI/Δ91*^mice as representative of the compound heterozygous phenotype in further studies.

To compare the extent of the prenylation defect in *Mvk* mutant mice and humans with MKD, we used freshly-isolated PBMCs from patients either homozygous for p.V377I or compound heterozygous for p.V377I and p.H20N variants. Importantly, the p.H20N mutation is in a highly conserved “hotspot” region (residues 8-35, around the active site of MK) where mutations are predicted to severely affect enzyme activity (Browne and Timson, 2015). We found that the pattern of defective protein prenylation in spleen cells from *Mvk* mutant mice bore striking similarity to that in PBMCs from humans with MKD. In other words, mice and humans with comparable genotypes had similar prenylation defects: mild in the homozygous p.V377I genotype and more pronounced in compound heterozygous genotypes bearing a bi-allelic combination of p.V377I with a more severe mutation (murine *Mvk*^*VI/Δ91*^ and human *MVK*^*VI/H20P*^) (Fig. 1E).

### *Mvk*^*VI/Δ91*^ mice have elevated mevalonic acid in plasma and cell extracts

Lack of MK activity causes the build-up of the substrate MA (Fig. 1D). Liquid chromatography-tandem mass spectrometry (LC-MS/MS) analysis of plasma revealed significantly higher levels of MA in *Mvk*^*VI/Δ91*^ compound heterozygous mice compared to wildtype *Mvk*^*+/+*^ animals. Plasma MA was similar between heterozygous *Mvk*^*+/VI*^, *Mvk*^*+/Δ91*^ and homozygous *Mvk*^*VI/VI*^ mice (Fig. 2A). Likewise, levels of intracellular MA were not different in bone marrow cell extracts from *Mvk*^*+/+*^, *Mvk*^*+/VI*^, *Mvk*^*+/Δ91*^ *or Mvk*^*VI/VI*^ animals but significantly higher in cell extracts from *Mvk*^*VI/Δ91*^ mice (Fig. 2B).

**Fig. 2.**
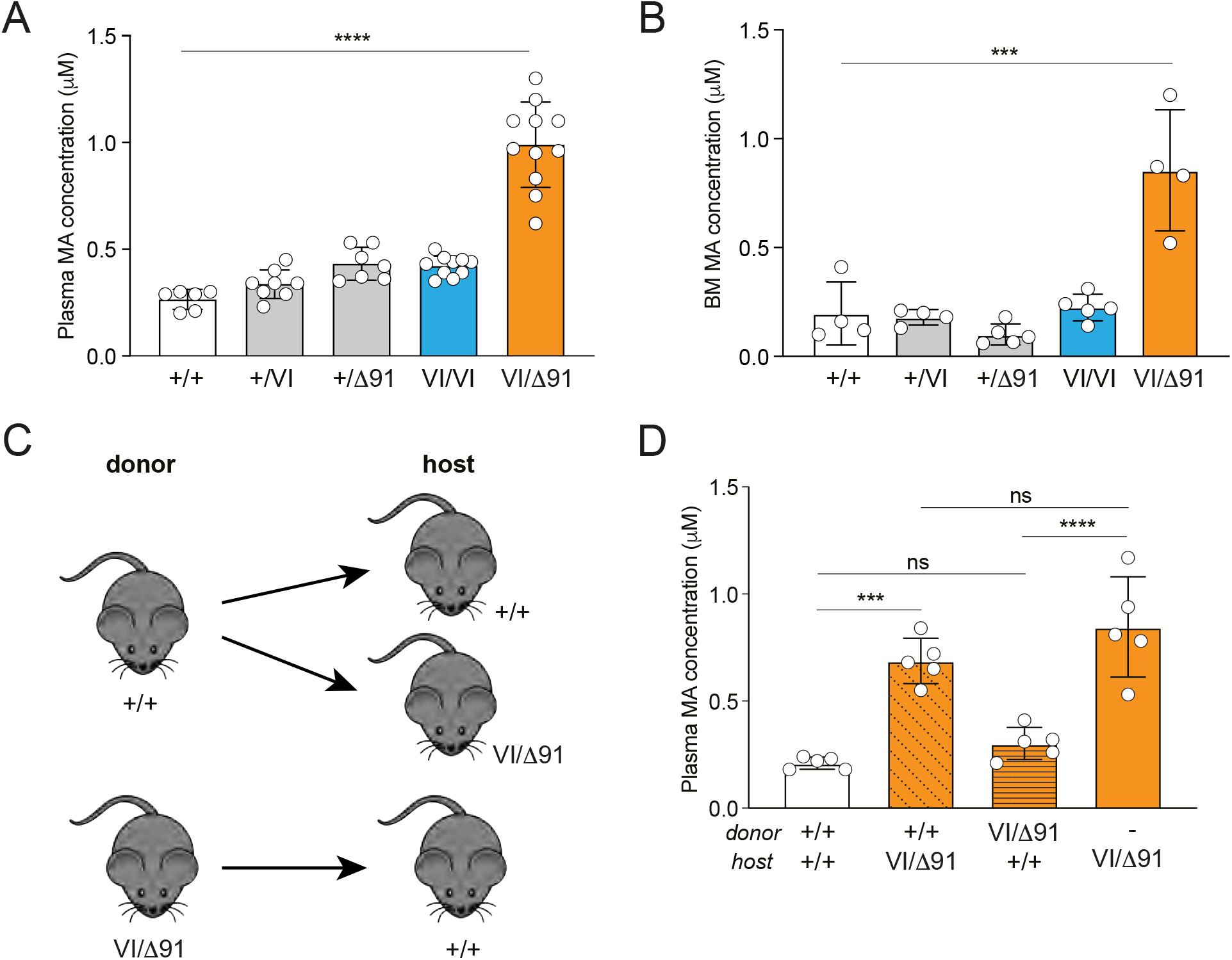
*Mvk*^*VI/Δ91*^ mice have elevated plasma mevalonic acid, from non-haematopoietic tissue. A) Concentration of mevalonic acid (MA) in plasma, or B) in bone marrow extracts, from wildtype and *Mvk* mutant mice. C) Scheme of bone marrow transfer from donor mice (wildtype or *Mvk*^*VI/Δ91*^) to generate chimaeric host mice. D) Concentration of MA in plasma from chimaeric mice and *Mvk*^*VI/Δ91*^ control mice. Values are mean ± SD (n = 6-11 mice per genotype in A, n=4-5 mice per group in B,D; each symbol represents a single mouse); ***p<0.001, ****p<0.0001, ANOVA with Tukey’s post-hoc test. Data in D) are representative of two independent experiments.

We used irradiated, chimaeric mice (Fig. 2C) to determine the contribution of the haematopoietic bone marrow cell compartment to the high levels of MA in plasma from *Mvk*^*VI/Δ91*^mice. Plasma MA levels remained low in wildtype hosts receiving *Mvk*^*VI/Δ91*^ mutant bone marrow, and comparable to non-chimeric wildtype controls (Fig. 2D) 8 weeks after bone marrow transfer. In contrast, *Mvk*^*VI/Δ91*^ recipients of wildtype bone marrow retained significantly elevated plasma MA, similar to non-chimaeric *Mvk*^*VI/Δ9*^ animals (Fig. 2D). Hence, whilst bone marrow cells may contribute a small amount to total plasma MA, non-haematopoietic tissues appear to be the major source of plasma MA in *Mvk*^*VI/Δ91*^ mice.

### p.V377I and Δ91 mutations affect a highly conserved core region of the MK protein

We used Aquaria (O’Donoghue et al. 2015; Kaur et al. 2021) to gain insights into how the *Mvk*^*VI*^ and *Mvk*^*Δ91*^ mutations affect MK enzyme activity. MK had a total of 119 related 3D structures, although none of these were exactly matching in sequence. The closest matches were two structures determined using rat MK (88% identical in amino acid sequence to mouse MK and an HHblits E-value of 3 × 10^−46^) (Steinegger et al. 2019). For further comparisons, we chose Protein Data Bank (PDB) entry 1kvk-A (Fu et al. 2008), since it also contains a bound ATP substrate.

Aquaria and the CATH (Class, Architecture, Topology and Homologous superfamily) structure classification database were used to identify relevant, evolutionarily conserved regions in MK (Sillitoe et al. 2021). We found two clear domains: 1) a C-terminal GHMP kinase domain (a family of kinases named after four of its members: galactokinase, homoserine kinase, mevalonate kinase, phosphomevalonate kinase; CATH superfamily 3.30.70.890) within residues 227-374; and 2) a N-terminal region spanning residues 7–226 homologous to a ribosomal protein S5 domain 2-type fold (also known as RPS5 domain 2; CATH superfamily 3.30.230.10) with an unusual three-residue segment close to the C-terminus (375–377). Importantly, Val377 is part of this interesting three residue region that folds back to form part of the N-terminal RPS5 domain (Fig. 3A; structure available [here](https://aquaria.app/Mouse/Mvk/1kvk?zenodo.3632187.V377I#(−11,-181,99))). Thus, the *Mvk*^*VI*^ (p.V377I) mutation would be expected to disrupt functions associated with this domain. Furthermore, the Val377 residue is buried from the solvent and forms part of the highly conserved, hydrophobic core of MK (Fig. 3B; structure available [here](https://aquaria.app/Mouse/Mvk/1kvk/A?zenodo.3632187.conservation#(−11,-181,99))). Any variants occurring in these core residues, even conservative substitutions as in p.V377I, are expected to be detrimental, and this is consistent with the partial loss of MK activity associated with the *Mvk*^*VI*^ allele. The remaining small segments on the N-terminus (1–6) and C-terminus (378–395) had no identifiable conserved homology CATH domain.

**Fig. 3.**
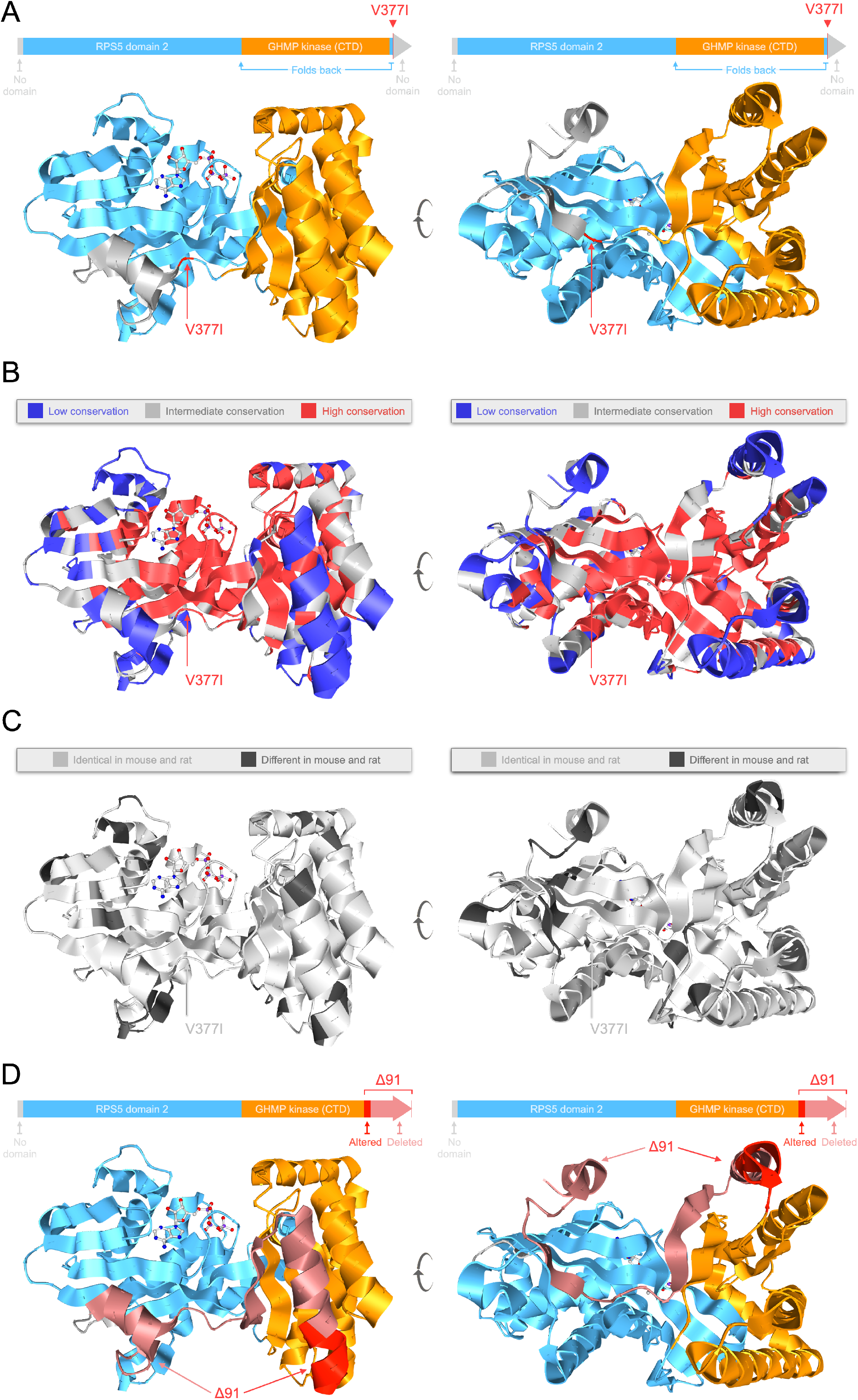
Visualisation of the conserved core region of mevalonate kinase affected by p.V377I and *Δ91* mutations. The figures show the crystal structure of a monomer of rat MK from PDB entry 1kvk-A, visualized using Aquaria. Left and right views are related by ∼90° rotation about the Y-axis. A) CATH domain assignments show the N-terminal RPS5 domain (blue), GHMP kinase domain (orange), and the position of the Val377 mutation within a 3-residue segment near the C-terminus that forms part of the RPS5 domain. B) Regions of low (blue), intermediate (grey) and high (red) sequence conservation, mapped in Aquaria using ConSurf conservation scores. C) Mouse-to-rat alignments from HHblits show that almost all residues in the conserved hydrophobic core (including the short segment around Val377) are identical (light grey) in mouse and rat. D) The *Δ91* mutation in *Mvk* alters residues 348-354 and causes premature truncation (deletion of all residues from 355 to the C-terminus, pink).

The amino acid sequences in the conserved hydrophobic core of MK are almost identical between mouse and rat (Fig. 3C; structure available [here](https://aquaria.app/Mouse/Mvk/1kvk/A?zenodo.3632187.mouse_vs_rat#(−11,-181,99))) and align with human MK without any gaps with the exception of a single residue inserted at the C-terminus: Leu-396. The *Mvk*^*191*^ mutation is predicted to alter residues 348-354 (LEQPEVE > PHTQLQL) and cause deletion of all remaining amino acids from 355 onwards. These changes involve the highly conserved core as well as both GHMP and RPS5 domains (Fig. 3D; structure available [here](https://aquaria.app/Mouse/Mvk/1kvk/A?zenodo.3632187.%CE%9491#(-11,-181,99))). Thus, the *Δ91* mutation would be expected to severely affect overall MK activity, and this is consistent with the complete loss of function associated with the *Mvk*^*Δ91*^ allele as described above.

### The effect of genetic disruption of *Mvk* differs from pharmacologic inhibition of the mevalonate pathway

Under steady state conditions, adult *Mvk*^*VI/Δ91*^ mice did not show any differences in the frequencies of B cells, T cells, dendritic cells, neutrophils or monocytes in peripheral blood, spleen or bone marrow (Suppl. Fig. 4A) compared to wildtype mice or *Mvk*^*+/VI*^ controls. Also, the level of inflammatory cytokines and chemokines in serum was barely detectable and did not differ between *Mvk*^*VI/Δ91*^ and *Mvk*^*+/VI*^ animals (Suppl. Fig. 4B). Similarly, serum IgD in wildtype and *Mvk* mutant mice was below the limit of detection of a commercial enzyme-linked immunosorbent assay (ELISA; data not shown). The lack of inflammation under basal/unstimulated conditions was striking when compared to a proposed pharmacologic model of MKD (Marcuzzi et al., 2013), which involves the *i*.*p*. administration of a bisphosphonate drug to acutely inhibit protein prenylation (Fig. 4A). Wildtype mice treated *i*.*p*. with the bisphosphonate alendronate (ALN), at comparable doses (6 mg/kg and 13 mg/kg) to those reported previously (Marcuzzi et al., 2013), showed a dramatic accumulation of uRab and uRap1A proteins in peritoneal cells 48 hours after treatment, a defect in prenylation that was far greater than in peritoneal cells from *Mvk*^*VI/Δ91*^ mice (Fig. 4B). Importantly, whilst the frequencies of immune cell populations in the peritoneal cavity of *Mvk* mutants were unchanged compared to controls (Fig. 4C,D), ALN treatment caused a dose-dependent infiltration of neutrophils, eosinophils and monocytes, and a striking decrease in large peritoneal macrophages (LPM) in wildtype animals (Fig. 4C,D). The higher dose of ALN also caused a clear reduction in the proportion of small peritoneal macrophages (SPM) and Ly6C^hi^ monocytes (Fig. 4C,D).

**Fig. 4.**
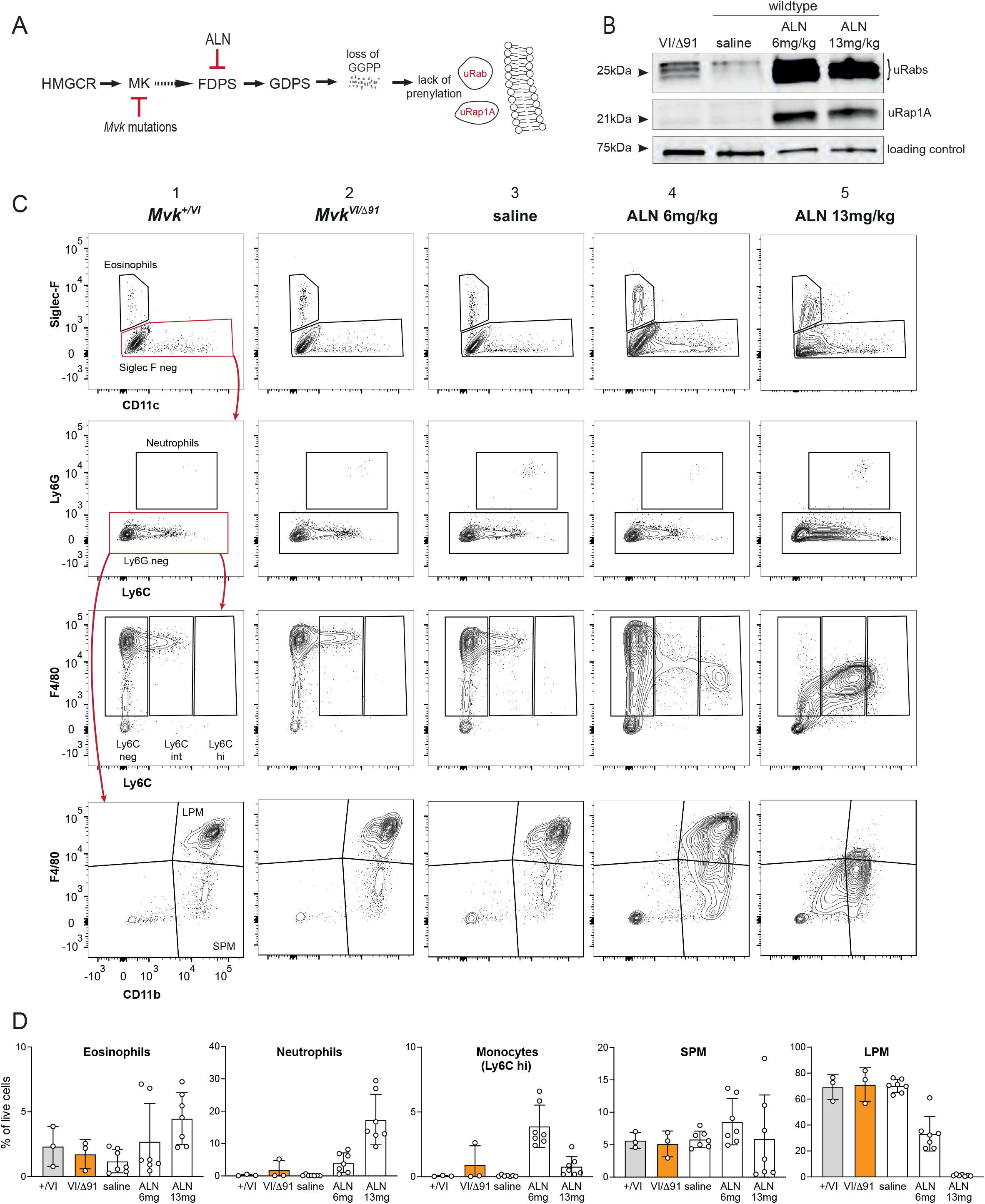
*Mvk*^*VI/Δ91*^ mice exhibit a lesser defect in prenylation and lack peritoneal inflammation compared with pharmacologic inhibition of the mevalonate pathway. A) Diagram of the mevalonate pathway and points of inhibition in *Mvk* mutant and in mice treated with the bisphosphonate drug alendronate (ALN). B) Analysis of unprenylated Rab GTPases (uRabs) and unprenylated Rap1A (uRap1A) in peritoneal cells from *Mvk*^*VI/Δ91*^ mice, and from wildtype mice 48 hours after *i*.*p*. treatment with alendronate (ALN: 6mg/kg or 13mg/kg) or saline control. C) Representative FACS plots of peritoneal cells. Each row illustrates the gating of eosinophils, neutrophils, monocytes, and large (LPM) and small (SPM) peritoneal macrophages. Columns 1-2 show FACS plots from *Mvk*^*+/VI*^ and *Mvk*^*VI/Δ91*^ mice, and columns 3-5 show FACS plots from wildtype mice treated with saline, 6mg/kg or 13mg/kg ALN. Polygons in red depict the population displayed in the proceeding plot (red arrow). D) Histograms show relative abundance (percentage of live cell singlets) of eosinophils (TCRb^-^, B220^-^, CD11c^-^, Siglec-F^+^); neutrophils (TCRb^-^, B220^-^, Siglec-F^-^, Ly6G^hi^); inflammatory monocytes (TCRb^-^, B220^-^, Siglec^-^F^-^, Ly6G^-^, F4/80^+^, Ly6C^hi^), LPM (TCRb^-^, B220^-^, Siglec^-^F^-^, Ly6G^-^, F4/80^hi^, CD11b^hi^) and SPM (TCRb^-^, B220^-^, Siglec^-^F^-^, Ly6G^-^, F4/80^+^, CD11b^+^). Data are the mean ± SD, each symbol represents a single mouse (n=3 per group for *Mvk*^*+/VI*^ and *Mvk*^*VI/Δ91*^mice, n=7 per group for ALN- and saline-treated mice).

### *Mvk*^*VI/Δ91*^ mice are hyper-responsive to endotoxin treatment and NLRP3 activation *in vivo*

PBMCs from *Mvk*^*VI/Δ91*^ mice had a clear defect in protein prenylation, with marked accumulation of uRab proteins compared to PBMCs from control *Mvk*^*+/VI*^ mice (Fig. 5A, Suppl. Fig. 3D). However, the proportion of circulating Ly6C^hi^ inflammatory monocytes was not significantly different between wildtype, *Mvk*^*+/VI*^ and *Mvk*^*VI/Δ91*^ mice (Fig. 5B), and freshly-isolated PBMCs from *Mvk*^*VI/Δ91*^ mice did not show any apparent difference in expression of a panel of 754 genes associated with myeloid innate immunity, compared to wildtype PBMCs (Suppl. Fig. 4C). Nonetheless, acute *in vivo* treatment of *Mvk*^*VI/Δ91*^ mice with *i*.*p*. LPS caused a significant increase in the levels of inflammatory serum cytokines and chemokines (IL-1β, IL-18, IL-6, G-CSF, IL-12, CCL2, CCL3 and CCL5) compared to control *Mvk*^*+/VI*^ mice (Fig. 5C,D). Several of these (G-CSF, IL-6, IL-12 and CCL2) were also significantly elevated in the peritoneal fluid of *Mvk*^*VI/Δ91*^ mice compared to *Mvk*^*+/VI*^ animals after *in vivo* LPS treatment (Suppl. Fig. 5A). These same inflammatory cytokines and chemokines (IL-1β, IL-18, IL-6, G-CSF, IL-12, CCL2, CCL3 and CCL5) were also higher in serum from a MKD patient, compared to healthy volunteers (Suppl. Fig. 6). In support of a role for NLRP3 in the enhanced inflammatory response *in vivo, i*.*p*. administration of a single dose of the NLRP3 inhibitor MCC950 (50mg/kg) 1 hour prior to LPS challenge completely abolished the LPS-induced increase in IL-18 and reduced IL-1β release to baseline serum levels in *Mvk*^*VI/Δ91*^ mice (Fig. 5E,F, Suppl. Fig. 5B). Pretreatment with MCC950 also significantly decreased serum IL-6 and CCL2 after LPS administration, and slightly reduced CCL5, but had no impact on G-CSF, CCL3 or IL-12 (Fig. 5F).

**Fig. 5.**
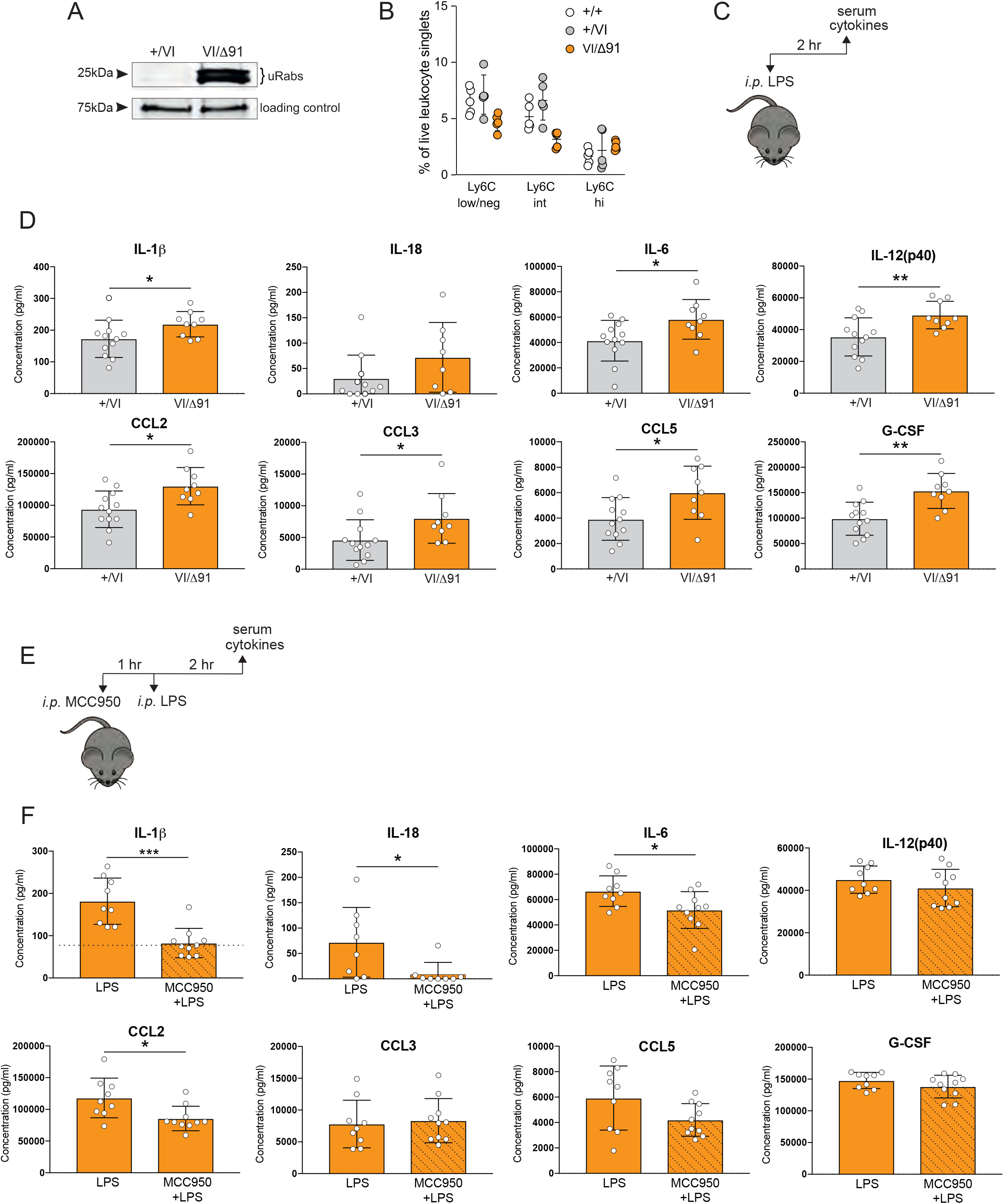
*Mvk*^*VI/Δ91*^ mice have elevated NLRP3-dependent IL-1β and other inflammatory mediators in serum following *in vivo* LPS treatment. A) Detection of unprenylated Rab GTPases (uRabs) in PBMCs from *Mvk*^*VI/Δ91*^ mice compared to *Mvk*^*+/VI*^ mice. B) Flow cytometric analysis of monocyte populations in PBMCs from wildtype, *Mvk*^*+/VI*^ and *Mvk*^*VI/Δ91*^ mice. Monocytes were gated as live leukocyte singlets negative for B220, TCRbeta, CD11c and Ly6G, and with low/negative, intermediate or high levels of Ly6C. Data are the mean ± SD (5 mice per genotype, each symbol represents a single mouse) and are representative of 3 separate experiments. C) *Mvk*^*+/VI*^ mice (n=12) and *Mvk*^*VI/Δ91*^ mice (n=9) were administered *i*.*p*. LPS 2 hours before serum collection. D) Serum cytokines and chemokines were measured using a multiplex assay and a separate IL-18 ELISA. E) *Mvk*^*VI/Δ91*^ mice were pre-treated with *i*.*p* 50mg/kg MCC950 1 hour prior to *i*.*p*. LPS challenge (n=9 with LPS alone, n=10 with MCC950 + LPS). F) cytokines and chemokines were measured in serum 2 hours after LPS challenge. The baseline level of serum IL-1β in untreated *Mvk*^*VI/Δ91*^ mice is shown by a dotted line (see also Suppl. Fig. 5B). Values are mean ± SD, each symbol represents a single mouse (*p<0.05, **p<0.01, ***p<0.001; unpaired t-test with Welch’s correction).

### Increased core body temperature exacerbates the defective mevalonate pathway *in vivo* in *Mvk*^*VI/Δ91*^ mice

Mutations in human MK protein may render the enzyme more sensitive to increased temperature (Houten et al., 2002; Mandey et al., 2006b), thereby potentially exacerbating the defect in protein prenylation in MKD. To test this hypothesis *in vivo*, we took advantage of the fact that homeostasis in mice becomes dysregulated at ambient temperatures above 30^°^C (Leon et al., 2010). To increase core body temperature (T_core_) we transferred cages of mice from the standard housing temperature (22 ± 1^°^C) to a heated chamber at 38^°^C for 18 hours (Fig. 6A). This procedure increased T_core_ in *Mvk*^*+/VI*^ and *Mvk*^*VI/Δ91*^ mice by 2.5^°^C (from approximately 34.5^°^C before heating to 37^°^C after heating; Fig. 6B) as measured using a digital rectal temperature probe.

**Fig. 6.**
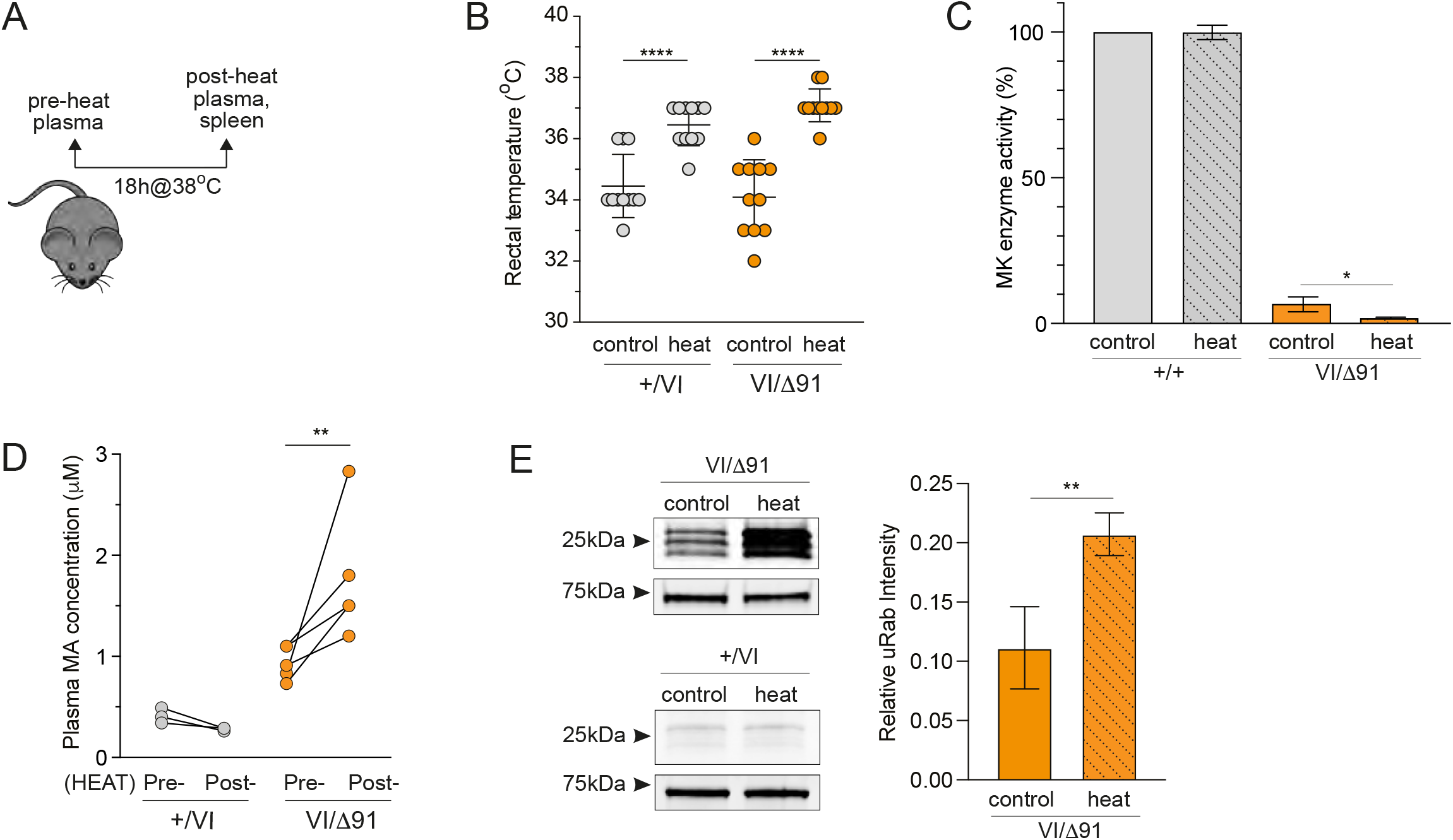
Elevated plasma mevalonic acid and defective protein prenylation are exacerbated *in vivo* in heated *Mvk*^*VI/Δ91*^ mice. A) *Mvk*^*+/VI*^ and *Mvk*^*VI/Δ91*^ mice were heated for 18 hours at 38^°^C. B) Rectal temperature pre- and post-heating. Bars show mean ± SD (n=11 mice per group, each symbol represents a single mouse; ****p<0.0001, ANOVA with Tukey’s post hoc test). C) MK activity in spleen cells from unheated and heated *Mvk*^*+/VI*^ and *Mvk*^*VI/Δ91*^ mice, expressed as a percentage of the MK activity in cells from an unheated *Mvk*^*+/+*^ mouse. Values are the mean ± SD, n=2 *Mvk*^*+/VI*^ mice and n=3 *Mvk*^*VI/Δ91*^ mice per group; *p<0.05, unpaired t-test. D) Levels of plasma MA pre- and post-heating (n=3 *Mvk*^*+/VI*^ mice, n=5 *Mvk*^*VI/Δ91*^ mice, each symbol represents a single mouse; **p<0.01, ANOVA with Tukey’s post hoc test). E) Unprenylated Rab GTPases (uRabs) in spleen cells from *Mvk*^*VI/Δ91*^and *Mvk*^*+/VI*^ heated and non-heated mice. For *Mvk*^*VI/Δ91*^ samples, blots were analysed by densitometry and values of uRab intensity were normalised to the loading control. Values are mean ± SD, n=4 mice; **p<0.01, unpaired t-test with Welch’s correction.

MK activity was 3-fold lower (and barely detectable) in spleen cells from heated *Mvk*^*VI/Δ91*^ compared to unheated mice, but was unaltered in spleen cells from heated and unheated control *Mvk*^*+/+*^ animals (Fig. 6C). Similarly, increased temperature did not alter the level of plasma MA in control *Mvk*^*+/+*^ mice but resulted in a significant rise in the already higher baseline level of plasma MA in *Mvk*^*VI/Δ91*^ mice (Fig. 6D). Furthermore, only heated *Mvk*^*VI/Δ91*^ mice had a clear exacerbation of the existing *in vivo* defect in Rab prenylation in splenocytes (Fig. 6E; Suppl. Fig. 7A). Despite the enhanced loss of prenylation in heated *Mvk*^*VI/Δ91*^ animals, there was still no indication (Suppl. Fig. 7B) of the immune cell infiltration and peritoneal inflammation observed in wildtype mice treated with bisphosphonate (Fig. 4).

### Prenylation in cells from mice and humans with MKD is sensitive to increased temperature

We further examined the effect of heat on *Mvk* mutant mouse cells by culturing freshly isolated bone marrow (BM) cells and bone marrow-derived macrophages (BMDM) at either 37^°^C or 40^°^C for 24 hours. Culturing at the higher temperature did not affect protein prenylation in wildtype BM (Fig. 7A). In contrast, the mild accumulation of uRabs in *Mvk*^*VI/VI*^ BM cells observed at 37^°^C was dramatically increased when cultured at 40^°^C (Fig. 7A).

**Fig. 7.**
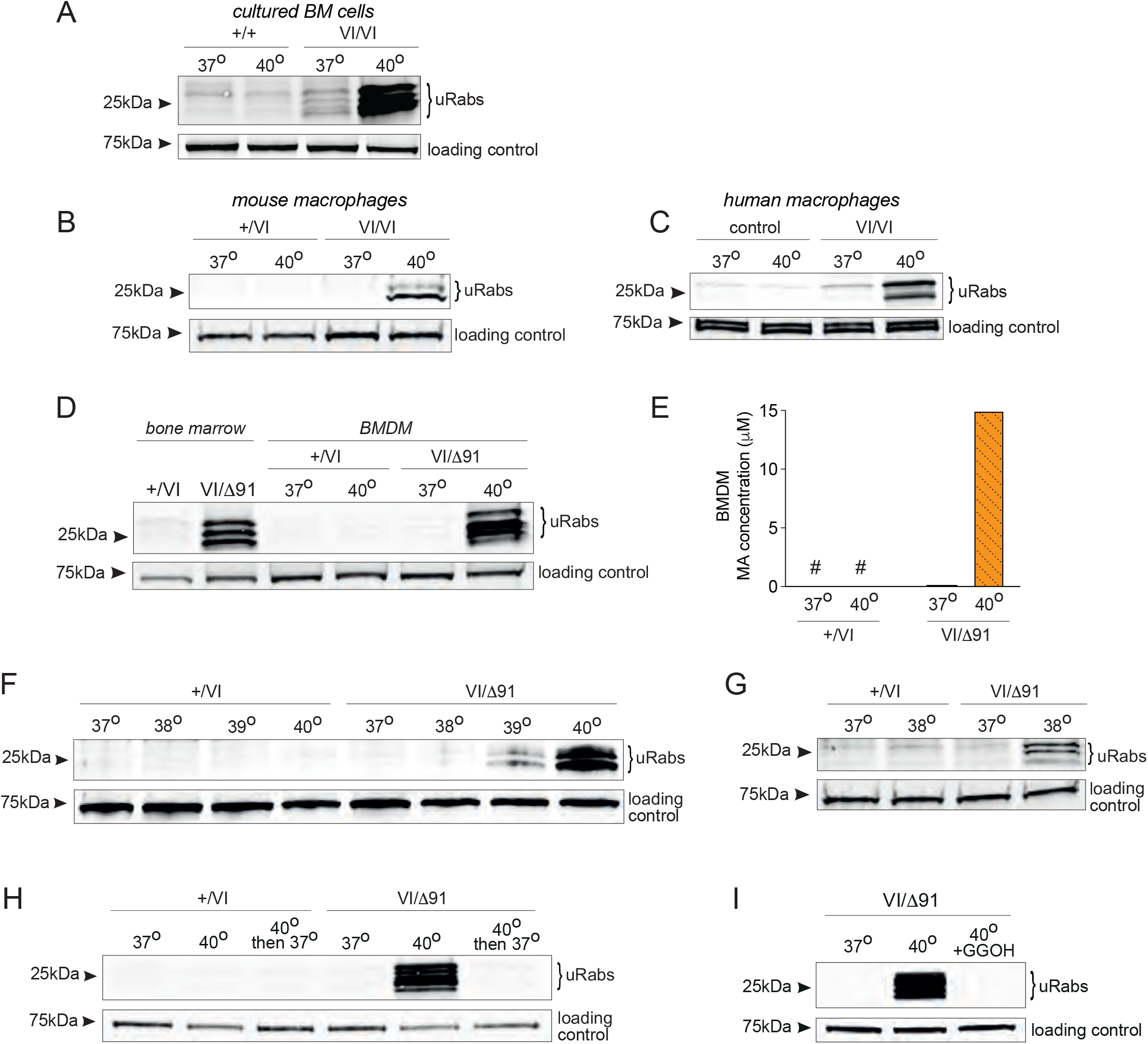
Increases in temperature enhance mevalonic acid accumulation and defective protein prenylation in cells from *Mvk* mutant mice and MKD patients. Analysis of unprenylated Rab GTPases (uRabs) in A) bone marrow cells from wildtype and *Mvk*^*VI/VI*^ mice cultured at 37^°^C or at 40^°^C for 24 hours; B) in BMDM from *Mvk*^*+/VI*^ and *Mvk*^*VI/VI*^ mice, cultured at 37^°^C or 40^°^C for 24 hours; C) in human monocyte-derived macrophages from a healthy control and *MVK*^*VI/VI*^ patient, cultured at 37^°^C or 40^°^C for 24 hours; D) fresh whole bone marrow and BMDM from *Mvk*^*+/VI*^ and *Mvk*^*VI/Δ91*^ mice, cultured at 37^°^C and 40^°^C for 24 hours. E) Concentration of MA in *Mvk*^*+/VI*^ and *Mvk*^*VI/Δ91*^ BMDM cell extracts after culturing cells at 37^°^C or 40^°^C for 24 hours (# = below limit of detection). Comparison of uRabs in *Mvk*^*+/VI*^ and *Mvk*^*VI/Δ91*^ BMDM cultured F) at 37^°^ to 40^°^C for 24 hours, G) at 37^°^ or 38^°^C for 5 days; and H) for 24 hours at 37^°^C, at 40^°^C, and at 40^°^C followed 24 hours recovery at 37^°^C. I) uRabs in *Mvk*^*VI/Δ91*^ BMDM cultured for 24 hours at 37^°^C or at 40^°^C in the absence or presence of 10μM GGOH. Blots shown in A,B,D,F-I are representative of 3 independent experiments.

Unlike fresh *Mvk*^*VI/VI*^ BM, *Mvk*^*VI/VI*^ macrophages lacked any detectable prenylation defect when grown at 37^°^C but re-gained uRab proteins when cultured at 40^°^C (Fig. 7B). A remarkably similar effect was observed when peripheral blood monocyte-derived macrophages from a HIDS patient of the same genotype (*MVK*^*VI/VI*^) were cultured at 40^°^C rather than at 37^°^C (Fig. 7C).

As with *Mvk*^*VI/VI*^ BMDM, defective protein prenylation was absent in *Mvk*^*VI/Δ91*^ macrophages compared to whole BM cells but re-appeared when cells were grown for 24 hours at 40^°^C (Fig. 7D). Elevated temperature (40^°^C) also caused a 50x increase in MA accumulation in *Mvk*^*VI/Δ91*^ BMDM but had no effect in control *Mvk*^*+/VI*^ BMDM (Fig. 7E). We also found that a 2^°^C temperature increase (39^°^C) for 24 hours was sufficient to cause the accumulation of uRabs in *Mvk*^*VI/Δ91*^ BMDM (Fig. 7F), whereas at 38^°^C a 5-day period was needed to have a detectable effect (Fig. 7G). Importantly, the dramatic buildup of uRab proteins in heated (40 ^°^C) *Mvk*^*VI/Δ91*^ BMDM was lost when cells were allowed to recover at 37^°^C for 24 hours (Fig 7H), presumably due to partial restoration of enzyme activity and degradation of unprenylated proteins that accumulated during heat exposure.

Increased temperature did not affect Rab prenylation in control *Mvk*^*+/VI*^ BMDM under any conditions (Fig 7B,D,F-H). Remarkably, the dramatic appearance of uRabs in heated *Mvk*^*VI/Δ91*^ BMDM was completely abolished in the presence of 10μM GGOH (Fig. 7I), an isoprenoid lipid precursor that can serve as substrate for protein prenylation when endogenous levels of GGPP are depleted.

## DISCUSSION

How defective flux through the mevalonate pathway leads to autoinflammation, and what mechanisms trigger inflammatory flares in MKD remain unclear, and this is largely due to the lack of genetic mouse models with *Mvk* mutations analogous to those present in MKD patients. We show here that mice bearing combinations of hypomorphic and amorphic mutations in *Mvk* recapitulate the characteristic biochemical features of HIDS, the milder form of MKD. Like humans heterozygous for pathogenic *MVK* variants, who have reduced MK activity (Faraci et al., 2021; Houten et al., 1999; Neven et al., 2007) but lack any detectable defect in protein prenylation (Munoz et al., 2017; Munoz et al., 2019), heterozygous *Mvk* mutant mice showed normal prenylation of small GTPases despite a 25-50% decrease in MK activity. Homozygous *Mvk*^*VI/VI*^ mice, harbouring the most frequent, hypomorphic HIDS missense mutation p.V377I (Boursier et al., 2021; Govindaraj et al., 2020; Houten et al., 2000; Ter Haar et al., 2016), had a mild prenylation phenotype that closely resembled the defect in PBMCs from a HIDS patient of the same genotype (*MVK*^*V377I/V377I*^) (Munoz et al., 2017; Munoz et al., 2019). In contrast, compound heterozygous *Mvk*^*VI/Δ91*^ mice completely lacking one functional *Mvk* allele had a more pronounced prenylation defect similar to that of a compound heterozygous *MVK*^*V377I/H20P*^ HIDS patient carrying the more severe p.H20P mutation (Munoz et al., 2017; Munoz et al., 2019).

Considering the extent of the defect in GTPase prenylation in *Mvk*^*VI/Δ91*^ and *Mvk*^*VI/VI*^ mice and their residual MK activity (9% and 19% respectively), these mouse avatars appear to represent opposite biochemical ends of the milder (HIDS) spectrum of MKD. Furthermore, by comparing the residual MK enzyme activity in heterozygous, homozygous and compound heterozygous mice with the extent of their prenylation defect, we found a distinct boundary around 20% residual MK activity. Above this boundary protein prenylation can be maintained normally (as is the case in *Mvk*^*+/Δ91*^ and *Mvk*^*+/VI*^ mice, with >40% MK activity), but below this 20% threshold insufficient synthesis of the lipid substrate GGPP leads to loss of protein prenylation (as in *Mvk*^*VI/Δ91*^ mice, with only 9% activity). This is entirely consistent with measurements of residual MK activities below 20% in compound heterozygous individuals with HIDS (Cuisset et al., 2001). *Mvk*^*VI/VI*^ mice appear to lie on this boundary, with 19% enzyme activity and a very mild prenylation phenotype. Humans homozygous for *MVK*^*V377I*^ may also sit on this precarious border, thus providing a new explanation for why some individuals homozygous for this variant are either unaffected or generally mildly symptomatic (Messer et al., 2016; Ter Haar et al., 2016). The ability of one *Mvk*^*VI*^ allele to confer about 10% residual MK activity *in vivo* in mice also explains why the *MVK*^*V377I*^ mutation is not associated with mevalonic aciduria in humans (Boursier et al., 2021; Brennenstuhl et al., 2021). In this most severe manifestation of MKD, residual MK activity is consistently <0.5% (Hoffmann et al., 1986; Hoffmann et al., 1993; Poll-The et al., 2000). We show, however, that one *Mvk*^*VI*^ allele confers ∼10% MK activity *in vivo* even if the second *Mvk* allele bears a complete loss of function mutation (as in *Mvk*^*VI/Δ91*^ mice), and this is enough residual activity to avert mevalonic aciduria. The reason why the conservative Val>Ile substitution at residue 377 reduces MK activity has remained puzzling, because this residue is distant from the active site (Browne and Timson, 2015). However, using Aquaria to visualize the 3D structure of MK we demonstrate that this residue lies within a short, highly conserved segment near the C-terminus (residues 375-377) that forms part of the hydrophobic core of the enzyme. In support of this, variants affecting the neighboring amino acid Gly376 have also been reported in patients with MKD in the online Infevers database (Sarrauste de Menthiere et al., 2003). Any alterations to this short segment, including p.V377I, are therefore likely to disrupt the critical core structure of the MK enzyme.

Our finding that the level of residual MK activity *in vivo* determines the extent of the defect in protein prenylation calls into question the physiological relevance of other proposed models of MKD in which the capacity for normal prenylation is far more severely compromised. These include, for instance, conditional deletion of geranylgeranyltransferase I (Akula et al., 2016), or acute inhibition of protein prenylation *in vivo* by *i*.*p*. administration of the bisphosphonate alendronate (Marcuzzi et al., 2008; Marcuzzi et al., 2013). We found that alendronate treatment in wildtype mice, at the two doses reported previously (Marcuzzi et al., 2008; Marcuzzi et al., 2013), had a far greater effect on prenylation in peritoneal cells than the endogenous defect in *Mvk*^*VI/Δ91*^ mice, but also caused a local inflammatory response that was absent in *Mvk* mutant mice. Alendronate-induced inflammation was clearly revealed by the influx of (Ly6Chi) monocytes, the increase in SPM and the loss of LPM in the peritoneal cavity after low dose treatment. The reduction in LPM frequency is consistent with inflammation-induced macrophage disappearance reaction, characterized by the migration of resident LPM into the omentum and increased adhesion to structural membranes (Liu et al., 2021). Increased cell death, a well-described effect of bisphosphonate drugs that inhibit prenylation (Luckman et al., 1998; Rogers et al., 2020), is the most likely cause of inflammation in the peritoneal cavity, and in particular the loss of both LPM and SPM/monocytes in mice treated with the higher dose of alendronate. The absence of detectable signs of inflammation in *Mvk*^*VI/Δ91*^ mice under steady state conditions (i.e. no change in baseline plasma cytokines/chemokines, unaltered immune cell frequencies) is consistent with the fact that HIDS patients are healthy between flares. Importantly, it also highlights the critical role of additional triggers that may tip the homeostatic balance towards exacerbating the biochemical deficiency in MK and/or repressing any metabolic compensation for lack of MK activity.

As well as recapitulating the prenylation phenotype of HIDS, the murine avatars of MKD described here also showed changes in plasma mevalonic acid (MA) that are consistent with defective MK activity. Elevated plasma and urinary MA is a characteristic feature of MKD first described in patients with mevalonic aciduria (Hoffmann et al., 1986; Hoffmann et al., 1993). However, urinary MA is also mildly raised in HIDS during inflammatory flares (Drenth et al., 1999; Houten et al., 1999; Houten et al., 2000; Poll-The et al., 2000). We found that, unlike the milder *Mvk*^*VI/VI*^ genotype, *Mvk*^*VI/Δ91*^ mice had slightly but significantly elevated MA in plasma and bone marrow cell extracts even under steady-state conditions, and this is consistent with their lower residual MK activity and greater defect in prenylation. *Mvk* mutant mice did not have elevated serum IgD. However, this feature is not considered to be a specific or reliable indication of HIDS/MKD in humans (Ammouri et al., 2007) and may be secondary to chronic inflammation (Di Rocco et al., 2001; Houten et al., 1999; Houten et al., 2000).

Bone marrow transplantation experiments revealed that the main source of MA in plasma was non-haemopoietic tissue, because the transfer of *Mvk*^*VI/Δ91*^ bone marrow into wildtype animals had little effect on the levels of circulating MA. Furthermore, plasma MA levels remained high in *Mvk*^*VI/Δ91*^ animals even after replacement with wildtype bone marrow. This is entirely consistent with reports that severe MKD patients that underwent successful haematopoietic stem cell transplantation (HSCT) maintained persistently elevated levels of urinary MA despite alleviation of fever and inflammatory symptoms (Faraci et al., 2021; Neven et al., 2007). Hence, urinary and/or plasma MA levels are not appropriate indicators of the success of HSCT therapy in MKD patients, reinforcing the need for careful monitoring of disease recurrence (Faraci et al., 2021). Liver, the main site of mevalonate-cholesterol biosynthesis (Goldstein and Brown, 1990), is likely the major source of the elevated plasma MA in mice and humans with MKD, at least under steady-state/non-febrile conditions. This also explains why treatment with simvastatin, a drug that selectively targets liver, decreased MA levels in HIDS patients (Simon et al., 2004a).

It has recently been suggested that MA can train innate immune cells to respond more robustly to stimulation via increased histone acetylation and epigenetic remodelling (Bekkering et al., 2018). We do not favour this as a physiological cause of inflammation in MKD because the concentration of MA used to train mouse macrophages *in vitro*, 500μM, (Bekkering et al., 2018) is 100x higher than the plasma level of MA we detected in *Mvk*^*VI/Δ91*^mice, and measurements of plasma MA even in mevalonic aciduria patients are mostly below 150μM (Hoffmann et al., 1993). Furthermore, statin therapy (which blocks MA synthesis) is considered not to be generally beneficial in the treatment of MKD (Lachmann, 2017), did not reduce the severity, frequency or occurrence of flares despite lowering MA urinary levels (Simon et al., 2004a), and even caused severe clinical crisis in two mevalonic aciduria patients (Hoffmann et al., 1993). Finally, MA levels remained persistently elevated in several MKD patients after HSCT despite the alleviation of fever and inflammatory symptoms (Faraci et al., 2021; Neven et al., 2007). Together, these observations strongly suggest that elevated MA is not the main underlying cause of inflammation in MKD although MA (or its derivative mevalonolactone) perhaps contributes to the neurological features of mevalonic aciduria (Cecatto et al., 2017). Indeed, neurological function in a patient with mevalonic aciduria substantially improved after liver transplant but inflammatory episodes only resolved after HSCT (Chaudhury et al., 2012).

Lack of normal protein prenylation is the most likely cause of inflammation in MKD. We recently provided the first direct evidence that protein prenylation is defective in freshly-isolated cells from individuals with MKD (Munoz et al., 2017; Munoz et al., 2019). Decreased prenylation (specifically, geranylgeranylation) of small GTPases such as Rac leads to enhanced inflammasome formation and subsequent increase in the processing and release of IL-1β (Jurczyluk et al., 2016; Kuijk et al., 2008a; Kuijk et al., 2008b; Mandey et al., 2006a; Skinner et al., 2019). However, how lack of prenylation leads to inflammasome activation, and the type of inflammasome that is activated, remain poorly understood and controversial, with evidence for a role of NLRP3 (Skinner et al., 2019), as well as pyrin ((Akula et al., 2016; Park et al., 2016; Skinner et al., 2019) inflammasomes. IL-1β release after LPS stimulation was elevated in *Mvk*^*VI/Δ91*^ mice compared to wildtype controls. This elevation was dependent on the NLRP3 inflammasome, since it was abolished by treatment with MCC950, a highly specific NLRP3 inhibitor (Coll et al., 2019; Coll et al., 2015). These findings are consistent with our recent report that the elevated IL-1β release from LPS-stimulated PBMCs from an MKD patient is abolished by MCC950 (Skinner et al., 2019), and suggest that inhibition of the NLRP3 inflammasome may be an appropriate therapeutic approach for MKD. However, consistent with evidence for increased production of a wide variety of proinflammatory cytokines from MKD PBMCs after stimulation (Stoffels et al., 2015), we also found significantly increased levels of serum IL-6, IL-12, G-CSF and CCL2, CCL3 and CCL5 in LPS-treated *Mvk*^*VI/Δ91*^ mice compared to controls, not all of which were reduced by MCC950 treatment. The inflammatory flares in MKD are therefore likely to be a complex, multi-cytokine-driven process. This is consistent with the fact that IL-1β-neutralising treatment is effective in some but not all MKD patients (Bodar et al., 2011; Jeyaratnam and Frenkel, 2020; Lachmann, 2017 9693), and therapies targeting IL-6 and TNF*α* can also be beneficial (Jeyaratnam and Frenkel, 2020).

Infection, vaccinations, strenuous exercise and psychological stress are recognised as frequent triggers of inflammatory flares in individuals with MKD (Drenth et al., 1994; Frenkel et al., 2001; Govindaraj et al., 2020; Houten et al., 2000; Lachmann, 2017). Notably, all of these can cause an increase in core body temperature (T_core_). Infection and vaccination can induce a PGE_2_-dependent rise in T_core_ (Morrison et al., 2014; Tupone et al., 2014). In a phenomenon known as psychological stress-induced hyperthermia (Oka, 2018), acute and chronic stress can also elicit an increase in T_core_ via an autonomic response that stimulates *Δ*3-adrenergic receptor-dependent thermogenesis in brown adipose tissue (Kataoka et al., 2020). Higher temperatures have been suggested to exacerbate the abnormal folding/stability of mutant MK protein (Browne and Timson, 2015; Houten et al., 2002; Houten et al., 1999; Houten et al., 2000; van der Meer, 2015). Consistent with this, and similar to MKD lymphoblast cell lines (Jurczyluk et al., 2016; Munoz et al., 2017), we found that *Mvk*^*VI/VI*^ and *Mvk*^*VI/Δ91*^ cells showed a striking sensitivity to elevated temperature. Even a 1^°^C rise in temperature for a few days was sufficient to develop a detectable defect in protein prenylation in cultured *Mvk*^*VI/Δ91*^ macrophages. A 3^°^C rise (equivalent to the high fever that is common in MKD patients (Frenkel et al., 2001) caused a dramatic build-up of unprenylated proteins in *Mvk*^*VI/Δ91*^ and *Mvk*^*VI/VI*^ cells after just 24 hours, an effect that was replicated with human *MVK*^*VI/VI*^ macrophages. The exacerbating effect of elevated temperature on defective prenylation was clearly reversible in *Mvk*^*VI/Δ91*^ BMDM when cells were returned to 37^°^C.

Most importantly, elevated temperature also worsened the metabolic defect *in vivo* in *Mvk*^*VI/Δ91*^ mice. Increasing T_core_ by 2-3^°^C further compromised the residual MK activity in splenocytes from 9% to <2%, and led to even higher levels of plasma MA and clear exacerbation of the defect in protein prenylation. This is reminiscent of the spike in urinary MA observed in HIDS patients during fever-associated flares (Drenth et al., 1999; Frenkel et al., 2001; Houten et al., 1999; Poll-The et al., 2000). Although the increase in plasma MA could reflect altered blood osmolality caused by dehydration in heated mice, this does not explain the dramatic buildup of intracellular MA in cultured *Mvk*^*VI/Δ91*^ macrophages after heating. Together, these findings provide compelling evidence that elevated T_core_ could trigger inflammatory flares in MKD by temporarily exacerbating the underlying defect in MK activity and protein prenylation, extending an idea proposed by Houten and colleagues 20 years ago (Houten et al., 2002). Approaches to minimise increases in T_core_, by pharmacological modulation of the central thermal regulatory network (Tupone et al., 2014), could therefore be considered as a future strategy to manage MKD. In addition, replenishment of isoprenoid lipid precursors (for example *via* dietary supplementation) should also be considered as an approach to prevent the worsening of defective prenylation in response to elevated T_core._ GGOH is a cell-permeable analogue of GGPP that can overcome the lack of isoprenoid lipid required for prenylation when upstream enzymes of the mevalonate pathway are inhibited pharmacologically, for example by a bisphosphonate (Fig. 4A) or statin (Benford et al., 1999; Skinner et al., 2019). We show that replenishing *Mvk*^*VI/Δ91*^ BMDM with GGOH completely prevented the dramatic intracellular buildup of uRabs in response to heat. *Mvk* mutant mice will therefore provide a much-needed preclinical platform to test the bioavailability, effectiveness and safety of GGOH or other lipid supplements as a preventative treatment in MKD.

The striking loss of the prenylation defect in cultured *Mvk*^*VI/VI*^ and *Mvk*^*VI/Δ91*^ BMDM compared to fresh bone marrow cells confirms reports that cells obtained or derived from MKD patients appear to adapt in culture (Houten et al., 2003). Furthermore, the reappearance of defective prenylation in heated BMDM is consistent with our previous observation that heat triggers loss of protein prenylation in MKD patient-derived cell lines (Jurczyluk et al., 2016; Munoz et al., 2017). This explains why abnormal prenylation has proven so difficult to demonstrate in cultured MKD cells (Houten et al., 2003), because they likely adapt by upregulating MK and other enzymes of the mevalonate pathway (Gibson et al., 1990; Hoffmann et al., 1997; Houten et al., 2003; Jurczyluk et al., 2016). Furthermore, infection of B lymphocytes with Epstein-Barr virus (EBV, which is routinely used to create immortalised lymphoblast cell lines from patients) causes upregulation of the mevalonate pathway (Wang et al., 2019). For this reason, accurate measurements of residual MK activity should ideally be performed with freshly-isolated PBMCs, rather than with cultured cells such as fibroblasts or EBV-transformed cell lines.

In summary, we show that novel mouse avatars of MKD, created by CRISPR/Cas9 editing of the *Mvk* gene, recapitulate the biochemical and clinical diagnostic features of the human disease. Our findings demonstrate a role of the NLRP3 inflammasome in the enhanced production of pro-inflammatory IL-1β and IL-18 *in vivo*. Furthermore, we highlight the likely role of increased T_core_ (for example in response to infection, stress or vaccination) as a mechanism for exacerbating the deficiency in the mevalonate pathway in MKD, thereby triggering inflammatory flares that may resolve once T_core_ returns to normal (Fig. 8). These new *in vivo* models of MKD will continue to shed further light on the pathophysiology of the disease and provide a valuable preclinical resource for testing new therapeutic approaches to overcome the deficiency in protein prenylation and/or prevent inflammatory flares.

**Fig. 8.**
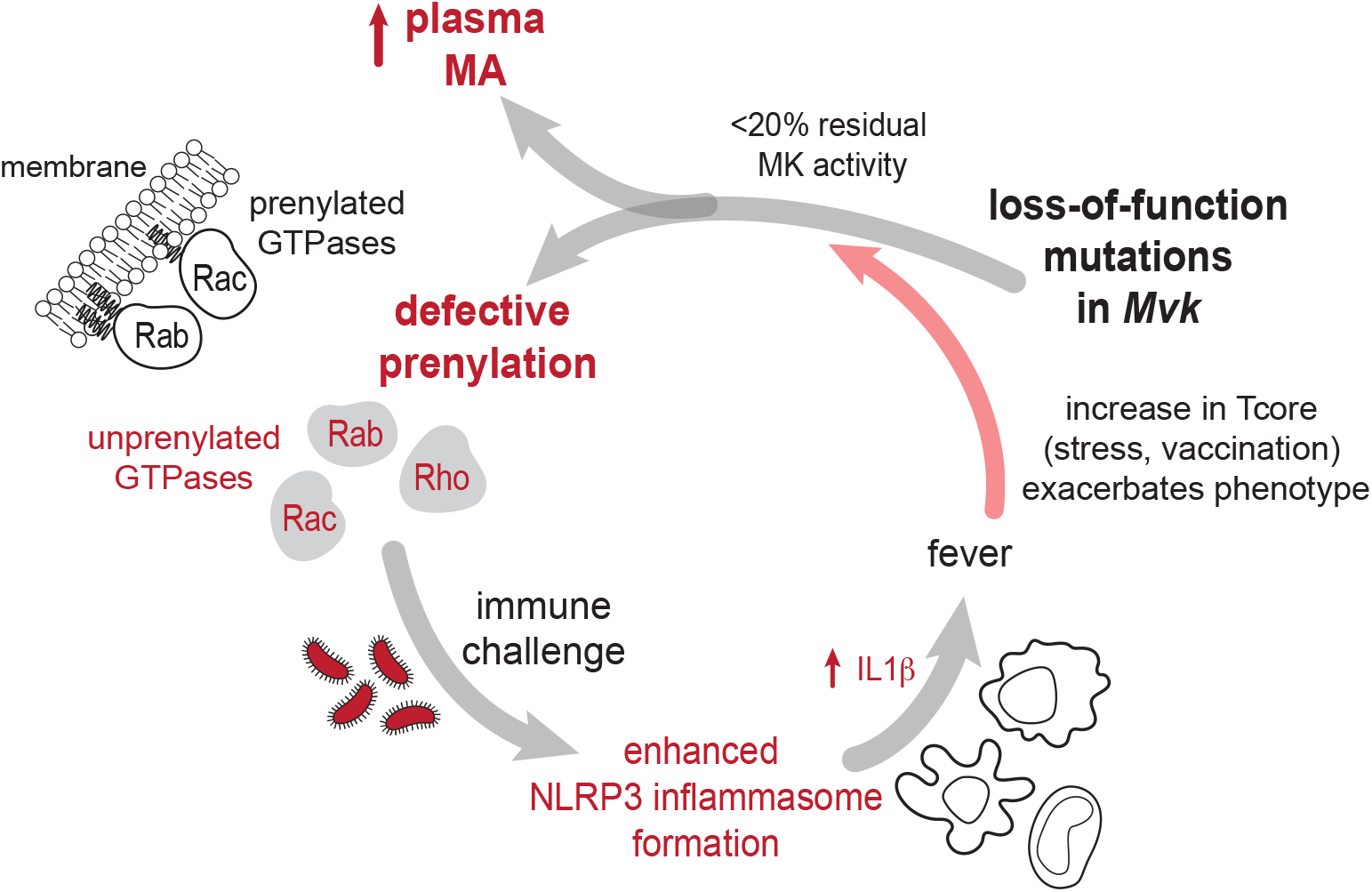
Pathophysiological features of inflammatory flares in MKD – lessons from mouse avatars. Genetic variants in *Mvk* causing <20% residual MK activity lead to a build-up of plasma mevalonic acid (probably derived mostly from liver) and intracellular accumulation of unprenylated small GTPase proteins. The latter enhance NLRP3 inflammasome formation after immune challenge, causing excessive release of IL-1β and other proinflammatory cytokines and chemokines. Elevated core body temperature (i.e. fever, vaccination or stress) exacerbates the defect in mutant MK enzyme, further worsening the metabolic deficiency in the mevalonate pathway and triggering an inflammatory flare.

## ACKNOWLEDGMENTS

We thank Prof Kirill Alexandrov and Dr Zakir Tnimov (University of Queensland) for providing reagents for the Rab prenylation assay, staff of the Garvan Institute’s MEGA, BTF and ABR facilities, Dr Yanchuan Shi, Tony Wang, Chris Jasieniecki (Garvan Institute), Isabelle Capell-Hattam and Prof Andrew Brown (UNSW Sydney), and Dr Rebecca Coll (University of Queensland) for advice and technical assistance. This work was supported in part by NHMRC project grant 1139644 to M.J.R., by grants to M.J.R. and M.A.M. from the St Vincent’s Clinic Foundation, the Allergy and Immunology Foundation of Australasia, the Marian & E.H. Flack Trust, the Mrs Janice Gibson and Ernest Heine Family Foundation, and by family members and friends of Maddison Dupond. O.P.S. and E.K.F. were supported through an Australian Government Research Training Program Scholarship. We also gratefully acknowledge funding by the New South Wales Government for the Victor Chang Cardiac Research Institute Innovation Centre, as well as funding from the Freedman Foundation for the Metabolomics Facility.

## AUTHOR CONTRIBUTIONS

M.A.M., M.J.R. and O.P.S. designed the experiments. M.A.M., O.P.S., J.J., Y.X., E.K.F., K.P. performed experiments and analysed data. E.M-F. performed flow cytometry analysis. E.K. and M.H. performed mass spectrometry analysis. D.Z. and R.B. performed CRISPR/Cas9 gene editing. E.D., A.A.B.R. assisted with experimental design. S.I.O’D. and S.K. performed the protein structure analysis in Aquaria. S.M., P.H., K.M-M., A.S. acquired patient samples. M.J.R. conceived ideas and oversaw the research program. M.J.R and M.A.M. wrote the manuscript.

## METHODS

### Generation of Mvk mutant mice

All experiments involving mice were approved by the Garvan Institute/St Vincent’s Hospital Animal Ethics Committee. *Mvk*^*V377I*^ and *Mvk*^Δ^ mice were produced by the Mouse Engineering Garvan/ABR (MEGA) Facility using CRISPR/Cas9 gene targeting in C57BL/6J mouse embryos following established molecular and animal husbandry techniques (Yang et al., 2014). The single guide RNA (sgRNA) was based on a target site in final exon (exon 11) of *Mvk* (AGCTGAGTGTGTGGAAACTC<u>CGG</u>, where CGG = protospacer-associated motif/PAM, underlined) and was microinjected into the nucleus and cytoplasm of C57BL/6J zygotes together with polyadenylated *S. pyogenes* Cas9 mRNA and a 140 base single-stranded, anti-sense, deoxy-oligonucleotide homologous recombination substrate carrying the p.Val377Ile (<u>G</u>TT><u>A</u>TT) mutation and a PAM-inactivating silent mutation in the P375 codon (CC<u>C</u>>CC<u>A</u>). A male founder mouse heterozygous for both substitutions was obtained and backcrossed with C57BL/6J female mice to establish the heterozygous *Mvk*^*V377I*^ line (referred to as *Mvk*^*+/VI*^). In addition, three mice carrying heterozygous frame shift deletions in exon 11 were bred to establish independent heterozygous *Mvk*^*Δ*^ lines. These lines carried 8bp, 13bp or 91bp deletions in exon 11 (*Mvk*^*+/Δ8*^, *Mvk*^*+/Δ13*^, *Mvk*^*+/Δ91*^) that resulted in frame shift mutations following the codons for T370, A374 and G347, respectively. Mice were crossed to generate homozygous *Mvk*^*VI/VI*^ mice and compound heterozygous *Mvk*^*VI/Δ8*^, *Mvk*^*VI/Δ13*^and *Mvk*^*VI/Δ91*^ mice. All mice were bred and housed with standard chow diet in specific pathogen-free conditions at Australian BioResources and the Garvan Institute Biological Testing Facility and used at 10-12 weeks of age.

### Bone marrow chimaeras

Recipient female *Mvk*^*VI/Δ91*^ (*Ptprc*^*b*^) and B6.SJL-*Ptprc*^*a*^*Pepc*^*b*^/BoyJArc mice (Australian Resources Centre), age 8-9 weeks, were irradiated twice with 425 Rads doses administered 6 hours apart. Irradiated mice were injected intravenously with 5-10 × 10^6^ bone marrow cells from female B6.SJL-*Ptprc*^*a*^*Pepc*^*b*^/BoyJArc or *Mvk*^*VI/Δ91*^ donor mice. 8 weeks later, plasma samples from chimaeric mice were analysed for mevalonic acid as described below.

### Immune cell isolation

Whole bone marrow was flushed from the femur and tibia and erythrocytes were lysed by incubating the suspension in lysis buffer (0.83% NH_4_Cl, 0.1% KHCO_3_) for 5 minutes at room temperature, and washed in cold Mg^2+^ and Ca^2+^-free DPBS (Gibco). Peripheral blood mononuclear cells (PBMCs) were isolated from age- and sex-matched mice by centrifugation over Ficoll-Paque Plus (GE Healthcare). Splenocytes were obtained by crushing the tissue through a 70μm nylon filter (Falcon), followed by erythrocyte lysis as above. Peritoneal cells were harvested by lavaging the peritoneal cavity with 5mL cold 2mM EDTA/ Mg^2+^ and Ca^2+^-free PBS (GIBCO). Isolated cells were then used to prepare cell lysates, for RNA extraction, or stained with conjugated antibody for flow cytometric analysis.

### PBMCs and monocyte-derived macrophages from MKD patients

Fresh samples of peripheral blood and serum were obtained from healthy volunteers and from MKD (HIDS) patients with confirmed pathogenic *MVK* variants – an adult male homozygous for p.V377I, an adult male compound heterozygous for p.V377I/p.Tyr149_Ser150insAlaTyr (Munoz et al., 2019), and a male child compound heterozygous for p.V377I/H20P (Munoz et al., 2017; Munoz et al., 2019). The study was approved by the Sydney Children’s Hospitals Network Human Research Ethics Committee (HREC/18/SCHN/403) and all subjects gave written informed consent in accordance with the Declaration of Helsinki. Buffy coat preparations of peripheral blood mononuclear cells (PBMCs) were isolated by centrifugation over Ficoll-Paque Plus (GE Healthcare). Cell pellets of PBMCs were snap frozen prior to analysis for unprenylated proteins.

To generate human PBMC-derived macrophages, frozen PBMCs from a 20-year old female with MKD (*MVK*^*Val377Ile/Val377Ile*^) and a healthy, age- and sex-matched volunteer were thawed and cultured in RPMI supplemented with 10% heat-inactivated fetal calf serum, 50 units/mL penicillin; 50μg/ml streptomycin (GIBCO), 100ng/ml rhM-CSF (Sino Biologicals) on bacterial plates in a humidified incubator (5% CO_2_) at 37^°^C for 7-10 days, replacing half the volume with fresh culture medium every 2 days. The resulting macrophages were then incubated either at 37^°^C or 40^°^C for 24 hours before collection. Cells pellets were analysed for the presence of unprenylated Rab proteins as described below.

### Bone marrow-derived macrophages

To generate bone marrow-derived macrophages (BMDM), cells were collected by flushing the femurs and tibias of adult mice with sterile PBS, and grown for 4 days on untreated tissue culture plates in a humidified incubator with 5% CO_2_ at 37^°^C with culture medium (RPMI, 10% heat-inactivated FCS, 50 units/mL penicillin; 50μg/ml streptomycin -GIBCO) supplemented with 50 ng/ml rhM-CSF (Sino Biologicals). The resulting adherent BMDM were removed by gentle scraping then re-plated and cultured for a further 24 hours at 37^°^C, 38^°^C, 39^°^C or 40^°^C, or at 38^°^C for 5 days, before harvesting. To test the effect of GGOH, a 10mM stock solution was prepared in ethanol and added to the culture medium at a final concentration of 10μM. Cell pellets were then analysed for the presence of unprenylated Rab proteins or mevalonic acid as described below.

### Detection of unprenylated proteins and western blotting

The accumulation of unprenylated proteins was assessed in murine and human tissue using an *in vitro* prenylation assay, as described previously (Ali et al., 2015; Jurczyluk et al., 2016; Munoz et al., 2017). Briefly, cell preparations or tissue samples were lysed by sonication in prenylation buffer (50mM HEPES, 50mM NaCl, 2mM MgCl_2_, 100µM GDP, 2mM DTT, 1x Roche complete EDTA-free protease inhibitor cocktail). 10-50μg of protein were incubated with recombinant GGTase II, REP-1 and biotin-GPP to label unprenylated Rab proteins with the biotinylated isoprenoid substrate (Ali et al., 2015). *In vitro*-prenylated (i.e. biotinylated) Rabs were then detected on PVDF membranes using streptavidin-680RD (LiCOR) (Ali et al., 2015). Some lysates were also analysed by western blotting for the presence of unprenylated Rap1A using goat anti-Rap1A (Santa Cruz, sc-1482) with anti-goat 680RD secondary antibody (LiCOR) (Ali et al., 2015; Rogers et al., 2020). β-actin (Cell Signalling, cat. #3700) or a narrow doublet of endogenous biotinylated protein approximately 73 kDa (often appearing as a broad singlet) were used as a sample loading control (Ali et al., 2015). Blots were scanned on a LI-COR Odyssey imager and analysed using Image Studio v5.2.5. MK protein was detected in liver homogenates by western blotting using 1μg/mL rabbit polyclonal antibody (Antibodies Online), raised to amino acids 179-228 of human MK, and HRP-conjugated anti-rabbit IgG. Blots were developed using SuperSignal West Pico reagent (Thermo Fisher) and scanned on a Fusion FX7 imaging system (Etablissements Vilber Lourmat SAS). Densitometry was performed using ImageJ (v2.0.0).

### LC-MS/MS analysis of mevalonic acid

For analysis of mevalonic acid in cells, 1mL cold extraction solvent (80:20 methanol:water) was added to frozen cell pellets then vortexed for 10 seconds and incubated in a ultrasonic bath filled with ice water for 1 hour, then centrifuged at 3000rpm for 30 minutes at 4^°^C. Aliquots (850μL) of supernatant were dried under vacuum in an Eppendorf Concentrator Plus then reconstituted in 42.5μL 70% methanol, 30% 10mM ammonium acetate. Samples (injection volume 1µL) were analysed by targeted LC-MS/MS using an Agilent 1290 Infinity II UHPLC system coupled to an Agilent 6495 triple quadrupole mass spectrometer. Separation was achieved using an Agilent Infinity Poroshell 120 EC-C18 column (3.0×150mm, 2.7µm) fitted with an Agilent Infinity Poroshell 120 EC-C18 UHPLC guard column (3.0×5mm, 2.7µm), maintained at 20^°^C. The mobile phases were 10mM ammonium acetate in water (A) and methanol (B), both containing 5µM medronic acid to chelate metal ions (gradient 98% A from 0 to 3mins, decreased to 2% A from 3.5 to 6.5mins at 0.5mL/mins, then increased to 98% A at 0.4mL/min from 6.5 to 12mins (total run time12mins). Autosampler temperature was 4^°^C. The mass spectrometer was operated in negative electrospray ionisation mode: source gas temperature was 250^°^C with flow at 17L/min, sheath gas temperature was 400^°^C with flow at 12L/min, and nebuliser pressure was 45 psi. Data were acquired in MRM (Multiple Reaction Monitoring) mode and was processed using Agilent MassHunter Quantitative Analysis software version B08.00.00. By comparison with a pure standard (Sigma Aldrich), mevalonic acid eluted at 1.9 min retention time. The limit of detection was 0.1μM.

### Measurement of MK activity

Immediately post-cull, liver was perfused with saline via the portal vein then snap-frozen. Slices of frozen liver, or frozen cell pellets of splenocytes and bone marrow prepared as described above, were homogenised in buffer (1:2.5 weight:volume for liver) containing 100mM KPO_4_ pH7.4, 5mM MgCl_2,_ 1mM DTT and 1x Roche complete EDTA-free protease inhibitor cocktail, using an ice-cold dounce homogeniser. After centrifuging (10,000*g* for 30 minutes, 4^°^C), MK activity in the supernatant was measured using a modification of previously-described methods (Gibson et al., 1989; Hoffmann et al., 1992). Briefly, 80ug of homogenate were added to a total volume of 23uL reaction buffer: 100mM KPO_4_ pH7.4, 6mM MgCl_2,_ 4mM ATP, with 0.33mM *R,S*-[2-^14^C]mevalonic acid (53mCi/mmol; Perkin Elmer) hydrolysed from the lactone form. After incubation at 37^°^C for 30 minutes, 1.7uL 88% formic acid were added then left for 30 minutes at room temperature. Following brief centrifugation to remove precipitate, 10uL of supernatant were spotted in duplicate onto Cellulose F thin layer chromatography plates (Millipore) and developed in n-butanol:formic acid:water (77:10:13) for 5 hours. Dried plates were exposed to a phosphorscreen and imaged using a FujiFilm Typhoon FLA5100 scanner. Mevalonate-5-phosphate and unreacted mevalonolactone were identified with R_f_ values 0.19 and 0.78 respectively, consistent with previous studies (Hoffmann et al., 1992). Densitometry to determine mevalonate-5-phosphate was performed using Multi Gauge v3.0 and MK activity was calculated as percentage of wildtype control.

### MK sequence and structure analysis

To find and examine all 3D structures related to mouse MK, we used Aquaria (O’Donoghue et al. 2015), which is based on sequence-to-structure alignments generated by HHblits (Steinegger et al. 2019). We also used the Aquaria interface to assign CATH domains (Sillitoe et al. 2021) and ConSurf conservation scores (Celniker et al. 2013), with the latter fetched from PredictProtein (Yachdav et al. 2014). PredictProtein/ConSurf assigned each residue to have low, intermediate, or high conservation, based on a Bayesian analysis (Mayrose et al. 2004) of evolutionary relatedness between the query protein and homologues in UniProt (The UniProt Consortium 2019). To create the figures showing structures mapped with domains, conservation, rat *vs* mouse sequence similarity, as well as the V377I and Δ91 variants, we used a version of Aquaria currently in development (O’Donoghue et al. 2021; Kaur et al. 2021) and available at https://aquaria.app.

### Alendronate treatment in vivo

A stock solution of 2.6mg/mL alendronate sodium salt (a kind gift of Dr FH Ebetino) was prepared by dissolving in saline, adjusted to pH 7.4. Mice (n=7 per group) were injected *i*.*p*. with 6mg/kg or 13mg/kg alendronate, or saline alone, then peritoneal cells were harvested 48 hours later by lavaging the peritoneal cavity with 5mL cold 2mM EDTA/ Mg^2+^ and Ca^2+^-free PBS (GIBCO). Cells were analysed by flow cytometry, or cells from 3 mice per group were pooled and analysed for unprenylated proteins, as described below.

### Flow cytometry

Mouse immune cell isolates from blood, bone marrow, spleen and peritoneal cavity were collected as described above and cell viability and total cell numbers were assessed by trypan-blue staining using a Corning CytoSMART cell counter. Cells were then pre-incubated with mouse-Fc block and viability dye (Zombie Aqua, BioLegend) in calcium/magnesium-free PBS (Gibco) for 15 minutes, before staining with fluorescently-conjugated antibodies (Supplementary Table 1) prepared in washing buffer (WB: 2mM EDTA, 0.02% azide; 0.5% foetal calf serum, calcium/magnesium-free PBS) or 45 minutes. Samples were rinsed 3 times in WB and fixed with 10% formalin neutral buffered solution (Sigma), washed and resuspended in WB for processing with a BD LSRII SORP flow cytometer/ DIVA software. The post-acquisition analysis was performed using FlowJo 10.6.2 (BD).

### Gene expression analysis

Blood was collected and pooled from wildtype or *Mvk*^*VI/Δ91*^ female mice (n=4 mice per genotype). PBMCs were isolated by centrifugation over Ficoll-Paque Plus (GE Healthcare) in SepMate™ tubes (STEMCELL Technologies, Canada) then RNA was extracted using TRIreagent (Sigma-Aldrich, T9424). 50ng RNA were hybridized at 65°C overnight according to the manufacturer’s instructions in a thermocycler using the nCounter Mouse Myeloid Innate Immunity Panel V2 (Nanostring Technologies, Seattle, WA). Hybridised samples were purified and immobilised onto a sample cartridge using the nCounter Prep Station. The cartridge was imaged and analysed using the nCounter Digital Analyzer and raw mRNA abundance frequencies were normalised to housekeeping and positive control genes using the nSolver Analysis Software 4.0. Values were expressed as log2 counts and plotted using ggplot2 R-package.

### Immune responses to LPS in vivo

12-week-old female *Mvk*^*VI/Δ91*^ and control *Mvk*^*+/VI*^ mice were injected *i*.*p*. with 100μg LPS endotoxin (*E*.*coli* O111:B4, Sigma-Aldrich) and culled 2 hours later for collection of serum and peritoneal fluid (the latter obtained by lavaging the peritoneal cavity of euthanized mice with 1mL PBS). In separate studies, *Mvk*^*VI/Δ91*^ mice were pre-treated with 50mg/kg *i*.*p*. MCC950 (or saline) 1 hour prior to *i*.*p*. LPS administration, then serum was collected 2 hours later. Baseline serum samples were prepared by bleeding mice 1 week before the experiment. Serum and peritoneal fluid were analysed for levels of cytokines and chemokines using a multiplex immunoassay (Bio-Plex, Bio-Rad) and a MAGPIX platform (Luminex), according to the manufacturer’s instructions. IL-18 was measured using a mouse IL-18 ELISA (Abcam).

### Elevation of core body temperature

Mice were bled by retro-orbital venepuncture to obtain a pre-treatment sample of plasma, and core body temperature (T_core_) measured 1 week later using a rectal probe (the whole procedure taking less than 10 seconds per animal to minimize stress). Mice were given a 1mL subcutaneous saline injection before heating for 18 hours at 38^°^C in a purpose-made chamber (2 mice per cage). T_core_ was again measured immediately after treatment, and post-treatment plasma samples and tissues collected. In some instances, mice were allowed to recover in normal housing conditions for 24 hours after heating. Mevalonic acid levels were analysed in plasma, and unprenylated Rab proteins were measured in spleen cells, as described above.

### Statistical analyses

Details of the statistical analyses performed are stated in the figure legends. Data are presented as the mean ± standard deviation (SD) and p values were calculated using unpaired two-tailed Student’s t test with Welch’s correction, or one-way ANOVA with Tukey’s post hoc test for multiple comparisons in GraphPad Prism v9. Significance was defined as *p < 0.05, **p < 0.01, ***p < 0.001, ****p < 0.0001 Alternatively, where stated in the figure legend, data shown are representative of at least 3 independent experiments.

