## Supplemental Figures and Legends for "Increased core body temperature exacerbates defective protein prenylation in mouse avatars of mevalonate kinase deficiency"

#### SUPPLEMENTARY FIGURE LEGENDS

**Supplementary Figure 1.** A) DNA sequence of exon 11 in wild-type and mutant mouse *Mvk* genes. The encoded amino acid sequence is indicated above the central base of the corresponding codons in the wild-type allele. The 20 base sequence corresponding to the Cas9-bound guide RNA (gRNA) is shown together with the associated protospacer-associated motif (PAM) and the central portion of the 140 base, single stranded oligionucleotide (ssOGN) homologous recombination (HR) substrate indicated below the wild-type allele. Targeted P375 and V377 codons are shown in ochre and green respectively, with altered bases in the V377I mutant highlighted in red. Inserted bases in the Δ8 mutant allele are indicated in blue. B) Sanger sequencing of wild-type *Mvk* exon 11 and mice heterozygous mice for the V377I, Δ8, Δ13 and Δ91 mutant alleles.

**Supplementary Figure 2. Western blot of mevalonate kinase in liver homogenate from wildtype and *Mvk* mutant mice**. Enhanced chemiluminescence detection with short exposure (left) and longer exposure (right) shows the full length, 40 kDa isoform of mevalonate kinase in wildtype (+/+) liver and *Mvk* mutant liver. The $Mvk^{VI}$ mutation does not detectably alter the mass of the major 42 kDa isoform of mevalonate kinase in $Mvk^{VI}$ or $Mvk^{VI/VI}$ liver. A truncated form, predicted to be generated from the $Mvk^{\Delta 91}$ mutation, or any degradation products could not be detected in $Mvk^{VI/\Delta 91}$ liver.

**Supplementary Figure 3. The mevalonate pathway is defective in bone marrow cells, spleen cells, PBMCs and peritoneal cells from homozygous and compound heterozygous *Mvk* mutant mice.** A) Detection of unprenylated Rab GTPases (uRabs) and unprenylated Rap1A (uRap1A) in bone marrow from Mvk mutant mice: homozygous $Mvk^{VI/VI}$ and three lines of compound heterozygous *Mvk* mutants compared to wildtype or heterozygous $Mvk^{+/VI}$ mice. B) Blots from prenylation assays of whole bone marrow samples were analysed by densitometry and values of uRabs normalised to the loading control (mean ± SD, n=3 mice per genotype). Values show wildtype (white), heterozygous (grey), homozygous $Mvk^{VI/VI}$ (blue), or compound heterozygous mice (orange). Detection and comparison of uRabs in C) spleen cells from $Mvk^{VI/\Delta 91}$ and $Mvk^{+/\Delta 91}$ mice, and D) in PBMCs from *Mvk* mutant mice (homozygous $Mvk^{VI/VI}$ and three lines of compound heterozygous *Mvk* mice), wildtype and heterozygous

$Mvk^{+/VI}$ mice. In panels A,C,D an endogenous, biotinylated 73 kDa protein was used as a loading control.

**Supplementary Figure 4. Baseline immune cell populations, gene expression in PBMCs and serum cytokine concentrations in $Mvk^{+/VI}$ and $Mvk^{VI/\Delta 91}$ mice.** A) B220$^{+}$ B cells, TCRβ$^{+}$ T cells, CD11c$^{+}$ dendritic cells (DC), B220$^{-}$ TCRβ$^{-}$ CD11c$^{-}$ CD11b$^{+}$ Ly6G$^{-}$ Ly6C$^{+}$ monocytes/macrophages (Mono/Mac) and B220$^{-}$ TCRβ$^{-}$ CD11c$^{-}$ CD11b$^{+}$ Ly6G$^{+}$ Ly6C$^{int}$ neutrophils (Neu), expressed as a percentage of live leukocytes in the bone marrow (left), blood (middle) or spleen (right) of $Mvk^{+/+}$ (no fill), $Mvk^{+/VI}$ (grey fill) or $Mvk^{VI/\Delta 91}$ (orange fill) mice (n=5 per group). Values are mean ± SD, differences were not statistically significant using a t-test corrected for multiple comparisons using the Holm-Sidak method. B) Cytokine and chemokine levels in serum samples from 12-week old female mice (n=11 $Mvk^{+/VI}$, n=8 $Mvk^{VI/\Delta 91}$). Cytokines were measured using a multiplex assay or a single cytokine ELISA for IL-18 (IL-18 levels were undetectable). Values are mean ± SD, ***p<0.001 (ANOVA with Tukey's post-hoc test); each symbol represents a single mouse. C) Gene expression analysis in freshly isolated PBMCs from $Mvk^{+/+}$ and $Mvk^{VI/\Delta 91}$ mice using an nCounter Myeloid Innate Immunity 754-gene panel (Nanostring). Cytokine/chemokine genes measured in (B) by multiplex immunoassay are highlighted. Red dotted line indicates no change in gene expression between $Mvk^{VI/\Delta 91}$ (y axis) and $Mvk^{+/+}$ (x axis), and grey dotted lines above and below indicate a log2 fold change of 1 or -1, respectively.

**Supplementary Figure 5. Elevation of inflammatory cytokines and chemokines in peritoneal fluid and serum of $Mvk^{VI/\Delta 91}$ mice following *in vivo* LPS treatment**. A) Peritoneal lavage fluid was collected from 12-week old female mice, 2 hours after *i.p.* injection of LPS (n=11 $Mvk^{+/VI}$, n=8 $Mvk^{VI/\Delta 91}$). Samples were analysed using a multiplex immunoassay. Values are mean ± SD (each symbol represents a single mouse), *p<0.05, **p<0.01, unpaired t-test with Welch's correction. B) Levels of IL-1β in serum samples from untreated (control) $Mvk^{VI/\Delta 91}$ mice (n=10), and $Mvk^{VI/\Delta 91}$ mice treated with *i.p.* LPS (n=9), or with 50mg/kg *i.p.* MCC950 prior to LPS treatment (n=10). Values are mean ± SD, ***p<0.001 (ANOVA with Tukey's post-hoc test); each symbol represents a single mouse.

**Supplementary Figure 6. Elevation of inflammatory serum cytokines and chemokines from an MKD patient compared to healthy controls**. Serum samples were from n=10 healthy controls, and from a MKD patient compound heterozygous for the pathogenic *MVK* variants p.V377I/p.Tyr149_Ser150insAlaTyr, previously described in Munoz *et al*, *Front Immunol* 10:1900 (2019). Cytokines and chemokines were measured using a multiplex immunoassay or a single cytokine ELISA for IL-18 (each symbol represents a single individual).

**Supplementary Figure 7. Elevated ambient temperature does not affect protein prenylation in wildtype animals or cause acute inflammation in $Mvk^{VI/\Delta 91}$ mice.** A) Analysis of unprenylated Rab GTPases (uRabs) in spleen cells from two wildtype (+/+) mice maintained at 22°C or two mice housed at 38°C for 18 hours (heated). An endogenous, biotinylated 73 kDa protein was used as the loading control. Spleen cells from a $Mvk^{VI/\Delta 91}$ mouse were used as a positive control on the same blot. B) Representative FACS plots from peritoneal cells isolated from untreated wildtype ($Mvk^{+/+}$) and $Mvk^{VI/\Delta 91}$ mice, and heated $Mvk^{VI/\Delta 91}$ mice. Polygons in red depict the populations displayed in the proceeding plot (red arrow). Histograms (right) show relative abundance (percentage of live cell singlets) from wildtype ($Mvk^{+/+}$), $Mvk^{VI/\Delta 91}$ and heated $Mvk^{VI/\Delta 91}$ mice. Immune cell populations were identified as follows: eosinophils ($TCRb^-$, $B220^-$, $CD11c^-$, Siglec-$F^+$); neutrophils ($TCRb^-$, $B220^-$, Siglec-$F^-$, $Ly6G^{hi}$); inflammatory monocytes ($TCRb^-$, $B220^-$, Siglec-$F^-$, $Ly6G^-$, $Ly6C^{hi}$), LPM ($TCRb^-$, $B220^-$, Siglec-$F^-$, $Ly6G^-$, $F4/80^{hi}$, $CD11b^{hi}$) and SPM ($TCRb^-$, $B220^-$, Siglec-$F^-$, $Ly6G^-$, $F4/80^+$, $CD11b^+$). Data are the mean ± SD, each symbol represents a single mouse (n=3 for wildtype and $Mvk^{VI/\Delta 91}$ mice, n=4 for heated $Mvk^{VI/\Delta 91}$ mice).

#### Supplementary Figure 1

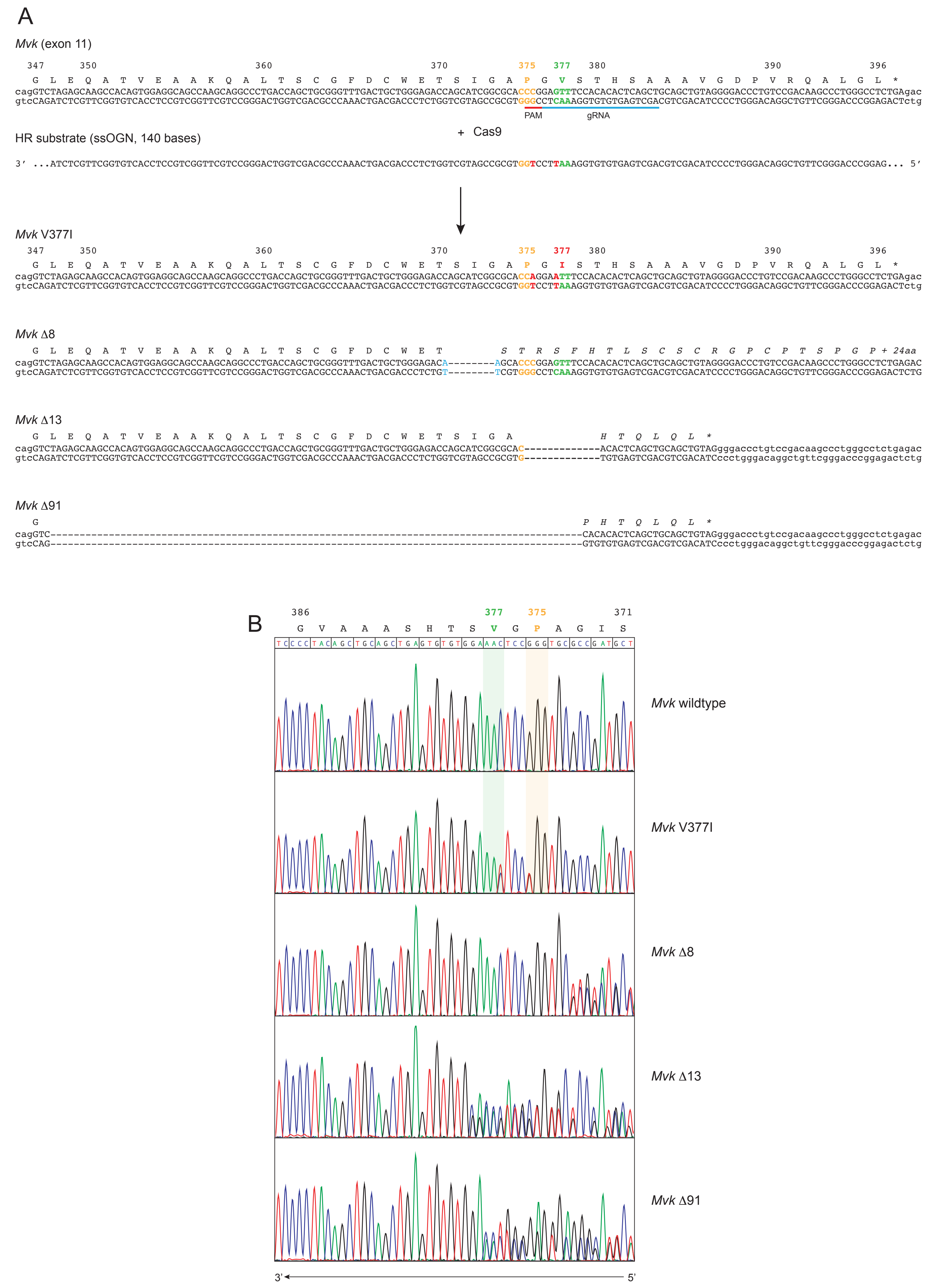

#### Supplementary Figure 2

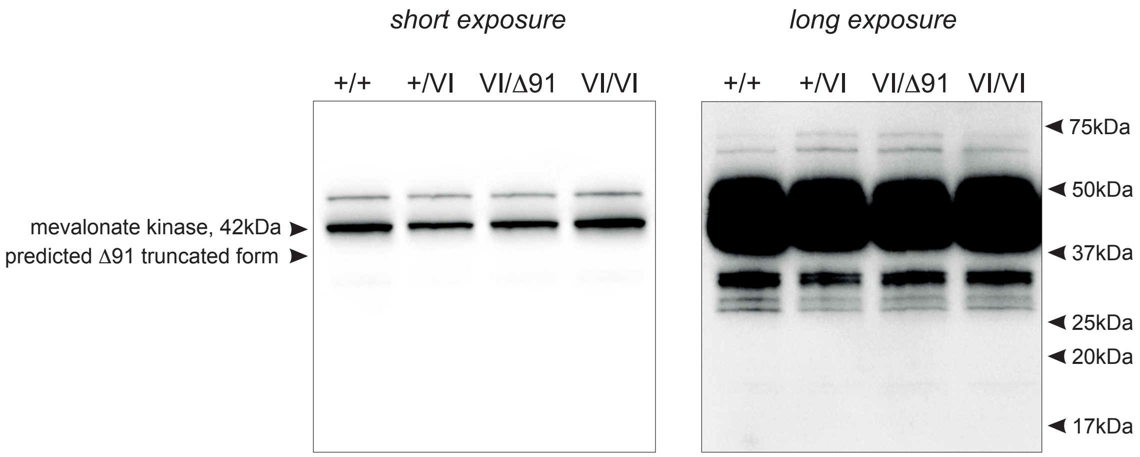

#### Supplementary Figure 3

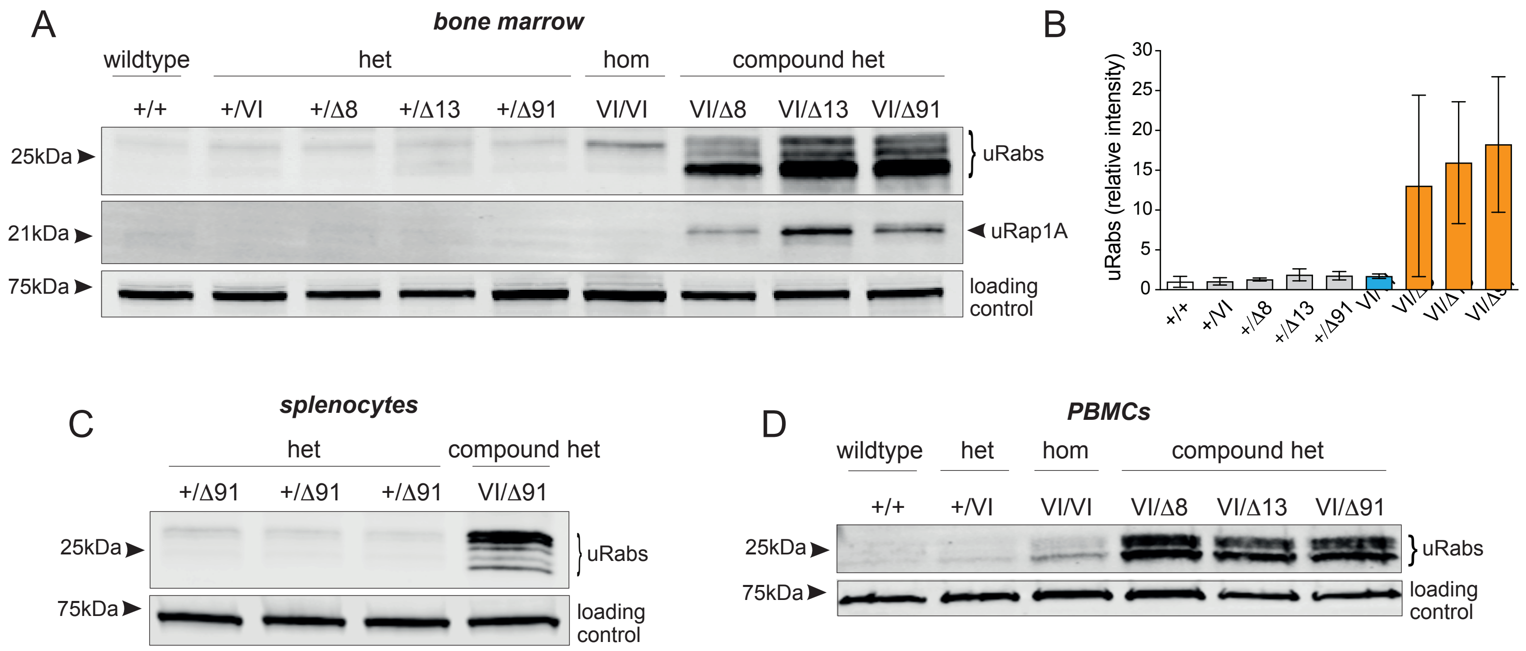

#### Supplementary Figure 4

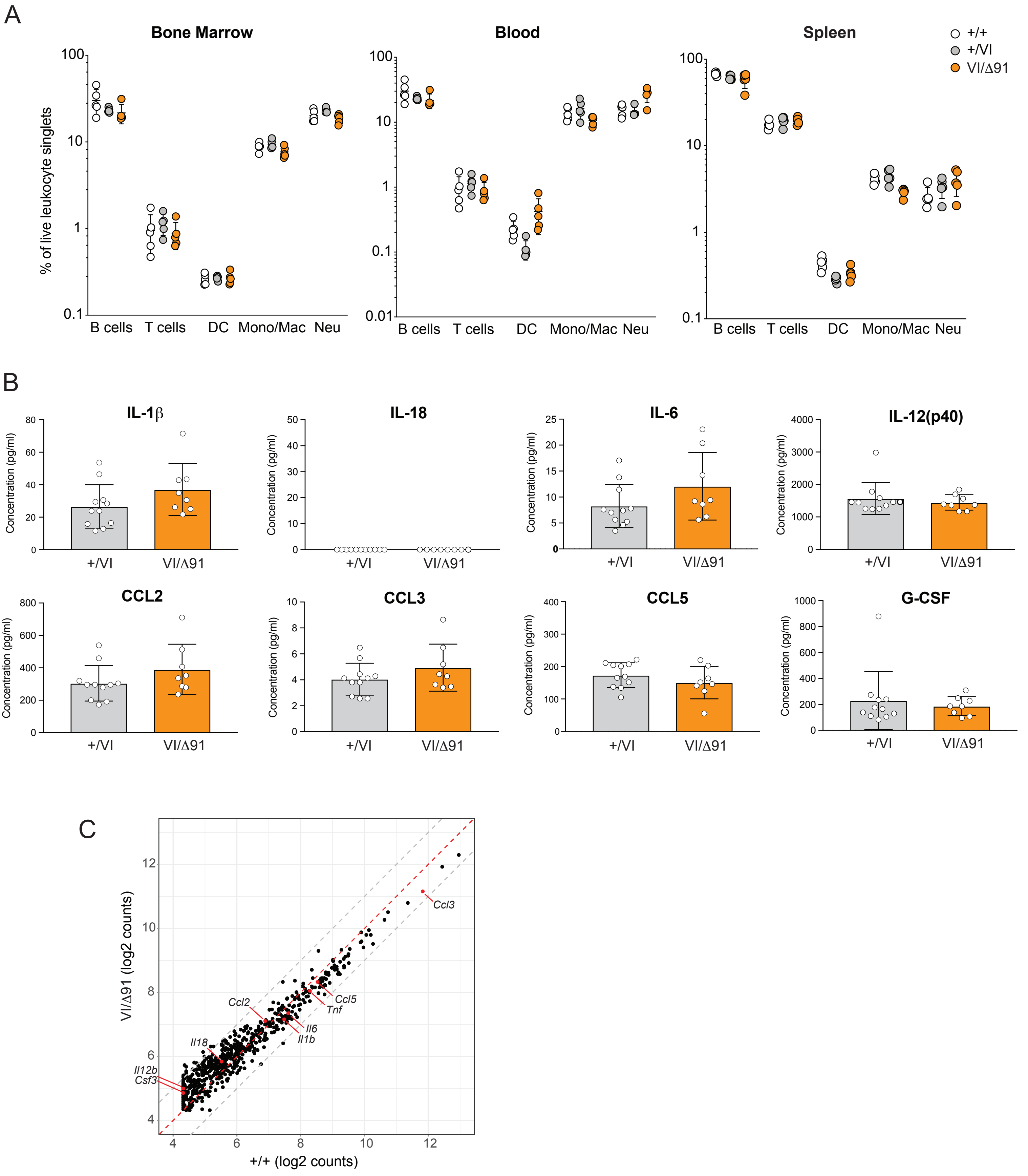

#### Supplementary Figure 5

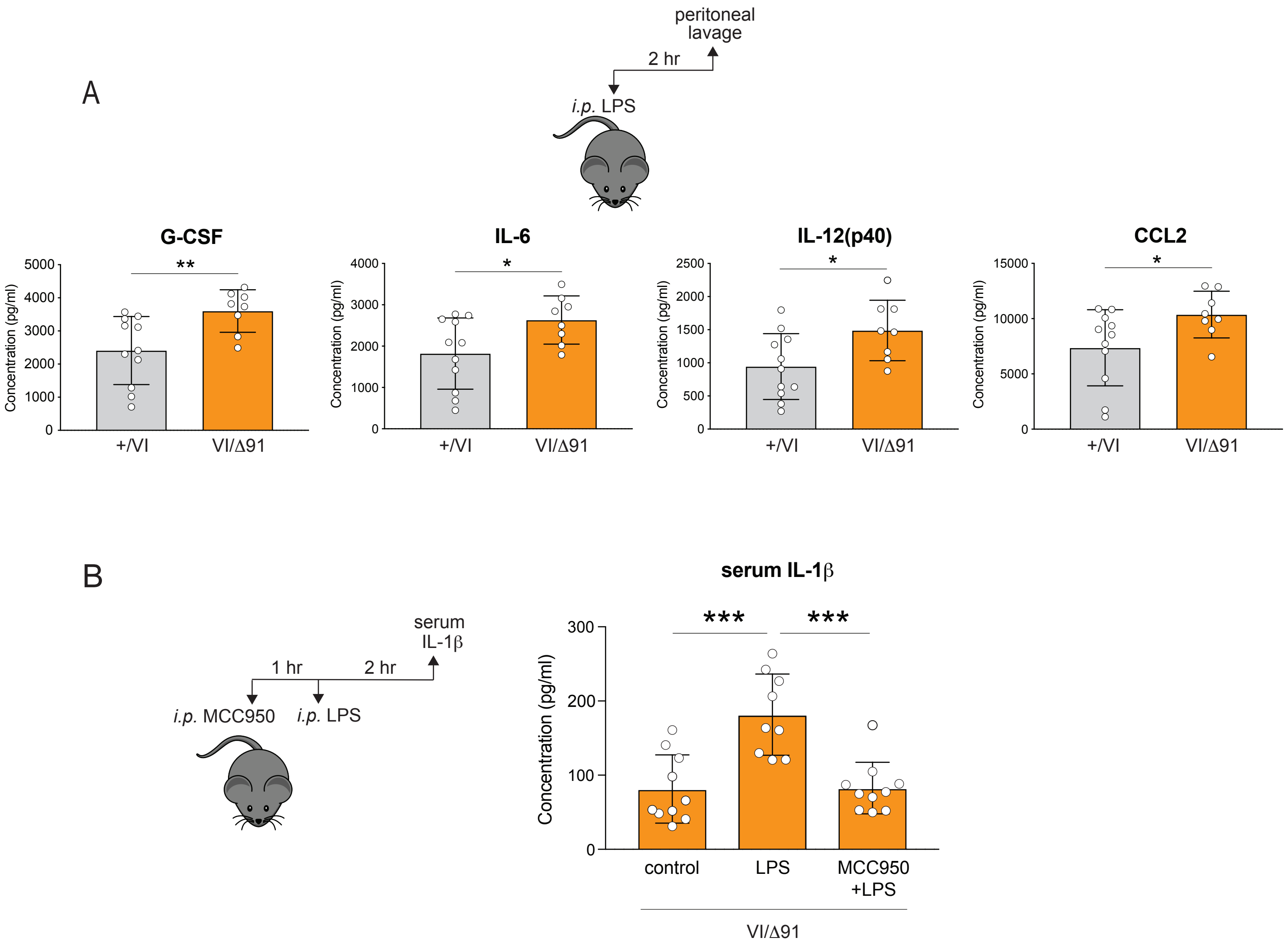

### Supplementary Figure 6

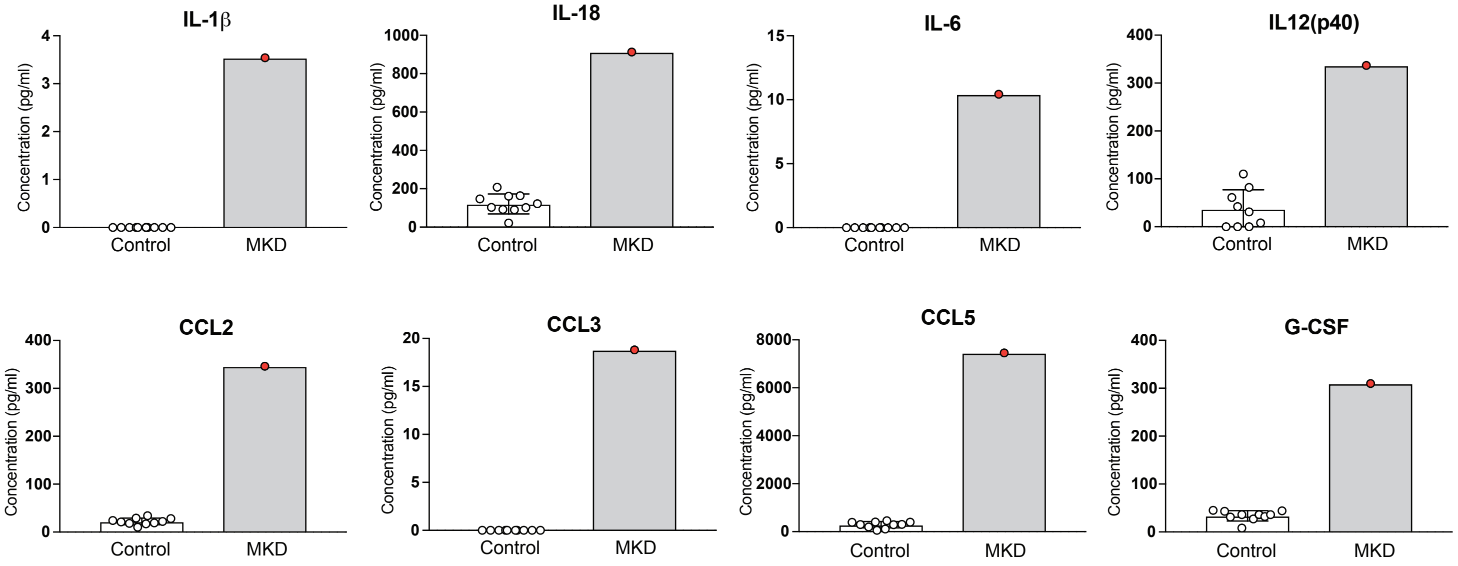

**Supplementary Figure 7**

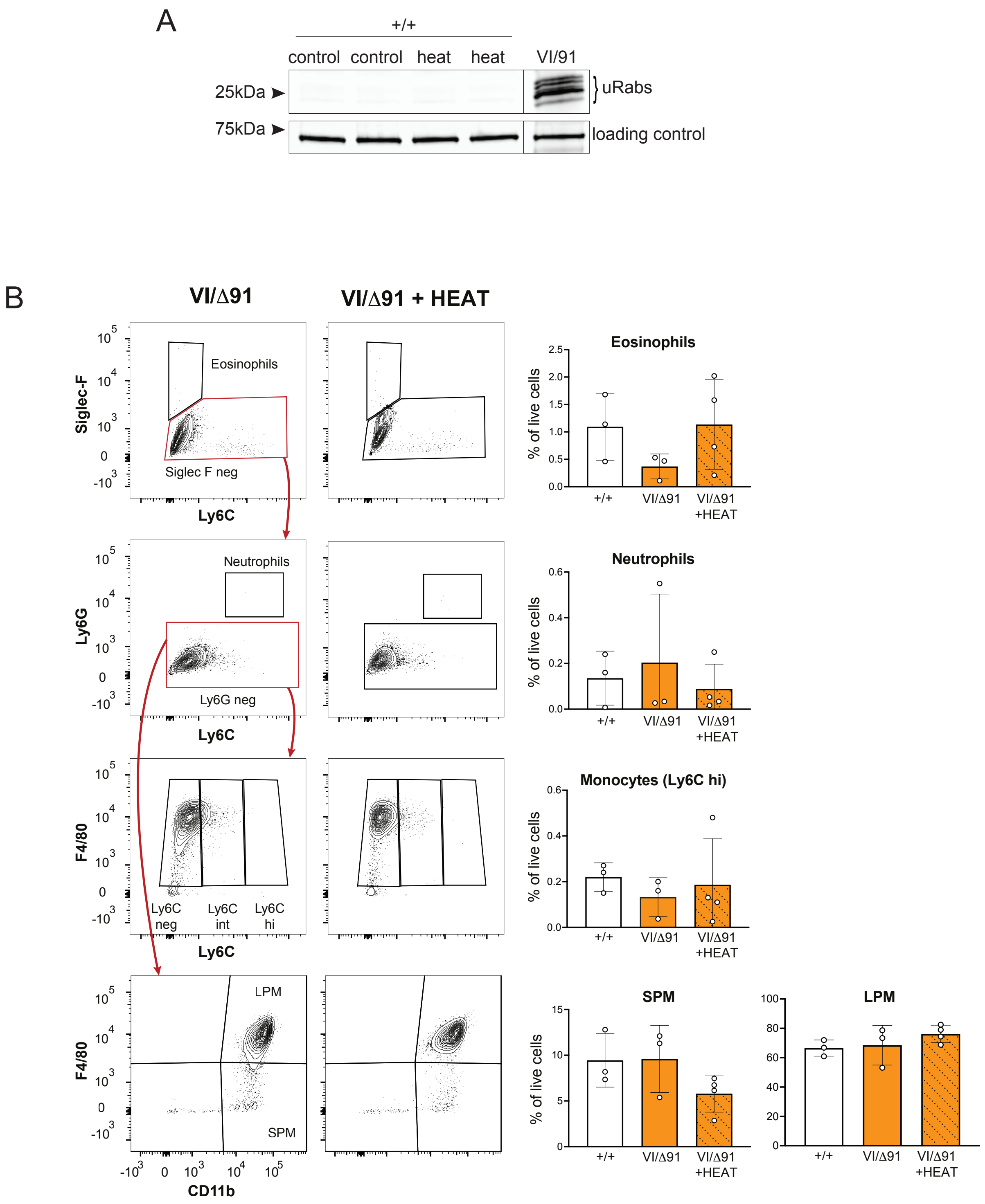
