## Supplemental Table 1 for "Increased core body temperature exacerbates defective protein prenylation in mouse avatars of mevalonate kinase deficiency"

**Supplementary Table 1.** Antibodies and reagents used for flow cytometry.

| Label | Target | Clone | Dilution | Supplier | Cat # |
|---|---|---|---|---|---|
| BUV395 | CD11b | M1/70 | 1/200 | BD Biosciences | 563553 |
| BV421 | F4/80 | T45-2342 | 1/100 | BD Biosciences | 565411 |
| BB515 | Siglec-F | E50-2440 | 1/200 | BD Biosciences | E50-2440 |
| PE | CD11c | N418 | 1/200 | eBiosciences | 117307 |
| APC | Ly6G | 1A8 | 1/200 | BD Biosciences | 560599 |
| BUV737 | B220 | RA3-6B2 | 1/300 | BD Biosciences | 612838 |
| PE-Cy7 | Ly6C | AL-21 | 1/200 | BD Biosciences | 560593 |
| APC-Cy7 | TCRB | H57-597 | 1/300 | BD Biosciences | 560656 |
| FITC | CD11b | M1/70 | 1/200 | BD Biosciences | 17-0051-81 |
| PerCP/Cy5.5 | CD11c | N418 | 1/200 | Thermo Fischer | 45-0114-82 |
| PE/Cy7 | Ly-6G | 1A8 | 1/200 | BD Biosciences | 560601 |
| Biotin | Ly-6C | AL-21 | 1/200 | BD Biosciences | 557359 |
| BV510 | CD19 | 6D5 | 1/100 | BioLegend | 115546 |
| BV786 | CD86 | GL1 | 1/100 | BD Biosciences | 740877 |
| BV605 | CD44 | IM7 | 1/300 | BD Biosciences | 563058 |
| PerCP/Cy5.5 | CD62L | MEL-14 | 1/200 | BD Biosciences | 560513 |
| Fc block | CD16/CD32 | 2.4G2 | 1/200 | BD Biosciences | 553142 |
| 7AAD | Dead cells | - | 1/500 | Invitrogen | A1310 |
| Zombie Aqua | Dead Cells | - | 1/500 | Biolegend | 423101 |
